# Operon-driven antagonism and novel modulation of DNA damage sensor STPK specific PP2C phosphatase in *Deinococcus radiodurans*

**DOI:** 10.64898/2026.01.23.701218

**Authors:** Ishu Soni, Dhirendra Kumar Sharma, Yogendra Singh Rajpurohit

**Author notes:** **Corresponding author and address** Dr. Yogendra Singh Rajpurohit 2-46-S, Modular Lab, A-block, Molecular Biology Division, Bhabha Atomic Research Centre Mumbai-400085 (Ex. 20387). **Conflict of Interest:** Authors have no conflict of interest in the content of the manuscript.

## Abstract

This study identifies a novel operon-driven signaling module in *Deinococcus radiodurans*. The operon includes a von Willebrand A domain protein (DRA0331), a Ser/Thr protein kinase (DRA0332), a canonical FHA-domain protein (DRA0333), and a PP2C-type phosphatase (DRA0334). DRA0334 shows Mn²□-dependent phosphatase activity and has a unique dual-domain structure that combines a Kinase-Interacting FHA (KI-FHA) domain with a PP2C catalytic domain. Functional assays show that FHA-domain protein, DRA0333 boosts the phosphorylation of STPKs like RqkA and DR1243 while operonic partner PP2C-type phosphatase, DRA0334 counteracts this through targeted dephosphorylation, establishing a phospho-regulatory antagonism. Notably, the KI-FHA domain of the DRA0334 phosphatase competitively interacts with the FHA domain to modulate the radiation-responsive RqkA kinase, thereby maintaining kinase–phosphatase balance. This KI-FHA domain also imparts substrate specificity and enables feedback regulation. Additionally, DRA0334 modular variants confirm separate roles of catalytic and docking modules, and STRING analyses link DRA0334 functions to DNA repair and stress recovery. Collectively, the findings suggest an operonic connection between DRA0333 and DRA0334, indicating that the KI-FHA and FHA domains may act as phospho-docking switches. These switches can regulate both kinase and phosphatase activities in a push–pull regulatory mechanism within the phosphorylation–dephosphorylation cycle of signal transduction, depending on their association with the type of catalytic domain.

## Introduction

Protein phosphorylation, catalyzed by serine/threonine/tyrosine protein kinases and reversed by cognate phosphatases, represents one of the most pervasive and evolutionarily conserved regulatory mechanisms governing cellular signaling, metabolism, and stress adaptation across all domains of life [1]. In bacteria, serine/threonine protein kinases (STPKs) and protein phosphatases (STPPs) create complex signaling circuits and regulate important processes such as cell division, morphogenesis, virulence, and stress recovery [1]. The balance between protein kinases and phosphatases controls phosphorylation levels at specific sites, influencing target proteins functions, localization, and interaction networking [1]. Although bacterial STPKs have been extensively studied, the mechanisms by which their cognate phosphatases fine-tune phosphorylation dynamics and substrate selectivity remain poorly understood. Increasing evidence suggests that phosphatase activity is not simply antagonistic to kinases but is intricately regulated by modular domains and protein–protein interactions that determine localization, substrate docking, and catalytic outcome [2].

In bacteria, STPKs and their related PP2C-type phosphatases regulate important bacterial activities like cell wall synthesis, autolysis, metabolism, cell division, and virulence. Examples include STPKs PknA/PknB in *Mycobacterium tuberculosis* [3], PrkC in *Bacillus subtilis*, Stk1 in *Staphylococcus aureus* [4], StkP in *Streptococcus pneumoniae* [5], and PknA/PknB/PknL in *Corynebacterium glutamicum* [6]. The specific and cognate Ser/Thr phosphatases, such as PrpC, PstP, Stp1, and PhpP, dephosphorylates different STPKs. For instance, PrpC works with PrkC from *B. subtilis* [7], PstP pairs with PknB from *M. tuberculosis* [8], Stp1 associates with Stk1 from *S. aureus* [9], and PhpP links to StkP from *S. pneumoniae* [10]. Thus, direct functional association of STPKs and STPPs reverse the STPKs induced signaling and regulate the signaling output of associated kinases.

In the setting of STPK and STPP coordinated signaling and functional interaction, phosphoprotein-binding modules offer another layer of regulation. Modules like SH2 and PTB recognize the phosphorylation of tyrosine residues within short linear motifs and facilitate assembling signaling complexes [11]. Contrary to tyrosine kinases, Ser/Thr kinases were initially thought to function primarily through phosphorylation-induced conformational changes and direct docking interactions in kinase-substrate proteins [12] but current knowledge suggested that modular phospho-binding domains specific to Ser/Thr phosphor-sites do exist. For example, 14-3-3 proteins and the WW domain of the proline isomerase Pin1 can bind directly to phosphoserine/phosphothreonine-containing motifs in their targets, forming dynamic signaling complexes [13]. Similarly, Forkhead-associated (FHA) domain was identified in a subset of Forkhead transcription factors [14] is a conserved phosphopeptide-binding module that specifically binds to phosphothreonine (pT) (Fig. S1 A). It is present in over 200 proteins found in both prokaryotes and eukaryotes.

Structurally, FHA domains are small, β-sandwich modules and that specifically recognize phosphorylated threonine (pThr) motifs within short sequence contexts and these modular domains come together to form multi-domain complexes, they are often called “super-domains,” highlighting their cooperative roles [15] (Fig. S1 B). The FHA domain typically consists of about 55–75 amino acids and features three highly conserved sequence motifs GR, SXXH, and NG separated by variable spacer regions [14]. Mechanistically, conserved arginine (GR) and serine (SXXH) residues recognize phosphothreonine, specificity comes from side chains in loop regions, which are often influenced by the amino acid at the pT+3 position (Fig. S1 A). In general, FHA domains achieve binding through rigid preformed surfaces. Minor adjustments in flexibility improve their affinity and specificity for phosphorylated targets [16].

In eukaryotes, the FHA domain plays a role in various processes, including signal transduction, transcription, DNA repair, protein degradation, and intracellular transport [15]. For instance, Rad53 from *Saccharomyces cerevisiae* has FHA1, for kinase recognition, and FHA2 for DNA damage checkpoint signaling [17], Fkh2 from *S. cerevisiae* combines FHA with a Forkhead transcription factor for gene regulation [18], Human tumor-suppressor CHK2 (hCHK2) kinase is homologue of Rad53 and FHA domain assist the oligomerization and autophosphorylation [19], KIF1A from *Mus musculus* combines a kinesin motor, FHA, and PH domain for intracellular transport [18], Nbs1 from *Homo sapiens* pairs FHA with a BRCT domain for DNA repair and checkpoint signaling [20], kinase-associated protein phosphatase (KAPP)-like PP2C phosphatase from *Arabidopsis thaliana* fuses FHA with a PP2C phosphatase to regulate plant receptor signalling [21]. Together, these examples highlight the functional diversity of FHA domain. In prokaryotes, FHA domains control a variety of processes, including secretion systems, sporulation, host-microbe interactions, and antibiotic resistance [22, 23]. Interestingly, the clustering of FHA-domain proteins with bacterial STPKs and STPPs has been noted in several bacteria including *Clostridium acetobutylicum*, *M. tuberculosis*, *Bacillus halodurans*, *D. radiodurans* and *Streptomyces coelicolor*, indicating key roles in coordinated molecular signaling.

*Deinococcus radiodurans* R1 stands out among bacteria for its extreme resistance to ionizing radiation and oxidative stress and has emerged as a valuable model to investigate stress-responsive signaling networks. This resistance comes from a combined efforts of strong DNA repair systems and effective antioxidant defenses [24-28]. Enzymatic antioxidants like catalases, peroxidases, and superoxide dismutases, along with nonenzymatic protectants such as pyrroloquinoline quinone (PQQ), carotenoids, and manganese complexes, shield both DNA and proteins from diverse reactive oxygen species (ROS) [29]. The DNA repair pathways in *D. radiodurans* are also remarkable, including nucleotide excision repair, base excision repair, mismatch repair, and specialized double-strand break repair via single-strand annealing (SSA) [30], extended synthesis-dependent strand annealing (ESDSA) [31], and their coordination is crucial for rebuilding the genome after significant DNA damage [32, 33]. Notably, *D. radiodurans* lacks error-prone pathways like translesion synthesis and nonhomologous end joining, which further supports its genetic stability by relying on homologous recombination [31].

The phosphorylation-mediated regulation by the Ser/Thr protein kinase RqkA is central to DNA damage induced signalling and influencing DNA repair and cell division progression [27, 34-42]. RqkA phosphorylates multiple cellular proteins including DNA repair protein (RecA, PprA) and key cell division related proteins (FtsZ, FtsK, and DivIVA), thereby controlling recombination and cell division during stress recovery. These modifications improve DNA repair while temporary halt in cell division, akin to eukaryotic DNA damage response (DDR) signaling by STPKs [43]. However, while phosphorylation-dependent activation of DNA repair and transient delay in cell division is well established, the corresponding dephosphorylation events catalyzed by Ser/Thr phosphorylation specific phosphatases and their regulatory control remain largely unexplored in *D. radiodurans* and not reported previously.

In the *D. radiodurans* genome, several STPKs exists and few genes encoding PP2C-type STPPs, forming putative kinase–phosphatase pairs reminiscent of eukaryotic signaling modules [44]. Among these, the DRA0331–DRA0334 operon presents a particularly intriguing organization. It encodes a von Willebrand type A (vWA) domain protein (DRA0331), a transmembrane STPK (DRA0332), a stand-alone FHA domain protein (DRA0333), and a PP2C-type phosphatase (DRA0334). This genomic clustering suggests tight functional coupling among these proteins, enabling coordinated regulation through a shared phosphorylation–dephosphorylation network that operates as an integrated signaling circuit rather than as isolated enzymatic events, thereby may ensure precise control of downstream cellular responses. Beside this, domain analysis of DRA0334 (PP2C-type phosphatase) reveals an unusual architecture: an N-terminal Kinase-Interacting FHA (KI-FHA) domain followed by a canonical PP2C phosphatase domain. The presence of an FHA domain within a phosphatase is rare in bacteria but is reminiscent of the kinase-associated protein phosphatase (KAPP) from *Arabidopsis thaliana*, which couples receptor-like kinase signaling to downstream regulatory feedback [21]. Such KI-FHA–PP2C combinations may represent evolutionary strategies for achieving phospho-dependent substrate recognition and feedback control. However, how the operonic KI-FHA and FHA modules interact at a mechanistic level and how this interaction shapes phosphorylation-mediated signaling remains unknown and is worth detailed reporting. This study uncovers a previously unrecognized operon-driven KI-FHA and FHA-domain antagonism in *D. radiodurans*, revealing that both kinases and phosphatases exploit FHA-mediated phosphosite recognition to regulate their activities. Research data suggest that these components constitute a modular, adaptable signaling unit that may help in fine-tunes phosphorylation homeostasis, potentially allowing *D. radiodurans* to rapidly balance kinase activation and phosphatase-driven reset possibly during DNA damage response and recovery.

## Results

### 1. DRA0331-DRA0334 operonic organisation and *D. radiodurans* Ser/Thr phosphatases analysis

Bacterial FHA (Forkhead-Associated) domain proteins are often found encoded by genes that cluster with genes encoding STPKs and STPPs [45] (Fig. S1 B). This genomic organization suggests functional relationships where the FHA domain proteins form a phosphoprotein interacting module for the precise docking, interact, and signalling modulation of target kinases and phosphatases [15]. In the Gram-positive bacterium *D. radiodurans*, a FHA-domain-containing protein, DRA0333, is located in a gene cluster with DRA0331-a von Willebrand type A domain protein, STPK-DRA0332, and STPP-DRA0334 (Fig. S2 A). This clustering in *D. radiodurans* indicates a potential functional phosphorylation/dephosphorylation link between STPK / STPP pair (DRA0332 / DRA0334) and the regulatory phosphor-adapter function of the FHA domain in DRA0333. Additionally, the existence of vWA-domain protein (DRA0331) in the operon might suggest the wider roles of FHA-domain mediated regulation of vWA domain protein DRA0331. vWA-domain proteins are highly phosphor proteins [46] and wide spread to play important roles in functions such as cell adhesion, protein-protein interactions, blood clotting, DNA repair and membrane transport [47]. Together, this observation supports the broader trend in bacteria where FHA domains frequently occur in close proximity to genes encoding the enzymes responsible for protein phosphorylation and dephosphorylation, suggesting that these components work together in bacterial signaling. The functional characterization of these clustered genes could reveal important aspects in *D. radiodurans*.

Furthermore, *in silico* analysis suggested that DRA0332 contains an STPK domain, spanning amino acids 50 to 314, and three transmembrane helices at its C-terminus while DRA0333 is a protein with a FHA domain, from amino acids 225 to 283, and DRA0334 features a PPM type STPP domain at the C-terminal, from amino acids 369 to 614 (Fig. S2 B). DRA0333 is an ortholog of cyanobacterial FraH protein and involved in possible oxidative stress and DNA repair as DRA0333 and its operon partner DRA0331 highly overexpressed in the novel oxidative stress sensor and transcription regulator (OxyR) mutant [48].

Comparative genomic analysis identified *dr2513* and *drA0334* genes encoding *D. radiodurans* specific PP2C phosphatases [44]. InterPro domain analysis (https://www.ebi.ac.uk/interpro) revealed that DR2513 contains an N-terminal PP2C domain spanning amino acids 6-242. Additionally, the TMhelix-2 server (https://services.healthtech.dtu.dk/services/TMHMM-2.0/) predicted two transmembrane regions at the C-terminal, located at amino acids 275-295 and 301-323. In DRA0334, the region encompassing amino acids 1-124 does not contain any functional domains and is classified as a disordered region while region from amino acids 210-368 contains a kinase interaction FHA-domain also designated as KI-FHA, while the C-terminal PP2C phosphatase domain is located between amino acids 369-614 (https://www.ebi.ac.uk/interpro/protein/UniProt/Q9RYH8/). The domain organization of DRA0334 resembles the kinase-associated protein phosphatase (KAPP)-like PP2C phosphatase of *A. thaliana* (refer AT5G19280_KAPP, Fig. S2 B), which interacts with various Ser/Thr kinases, such as RLKs, SERK1, CLV1, and FSL2 in *A. thaliana* and other plant species, playing a role in meristem development [21]. The sequence of FHA-domain of DRA0333 (amino acids 225-283) has typical FHA-domain specific motifs like GR, SXXH, and NG, however KI-FHA domain of DRA0334 showed less conservation for the typical FHA domain specific motifs suggested that KI-FHA domain has either different phosphor-residue specificity than phosphor-threonine residue for which typical FHA domain has recognition sites (Fig. 1A). Similar type of less conserved FHA domains have been shown for several proteins such as all0157, SC6D10.12, alr1603, CT664, and orf/myxxa/FHA2 [45].

**Figure 1.**
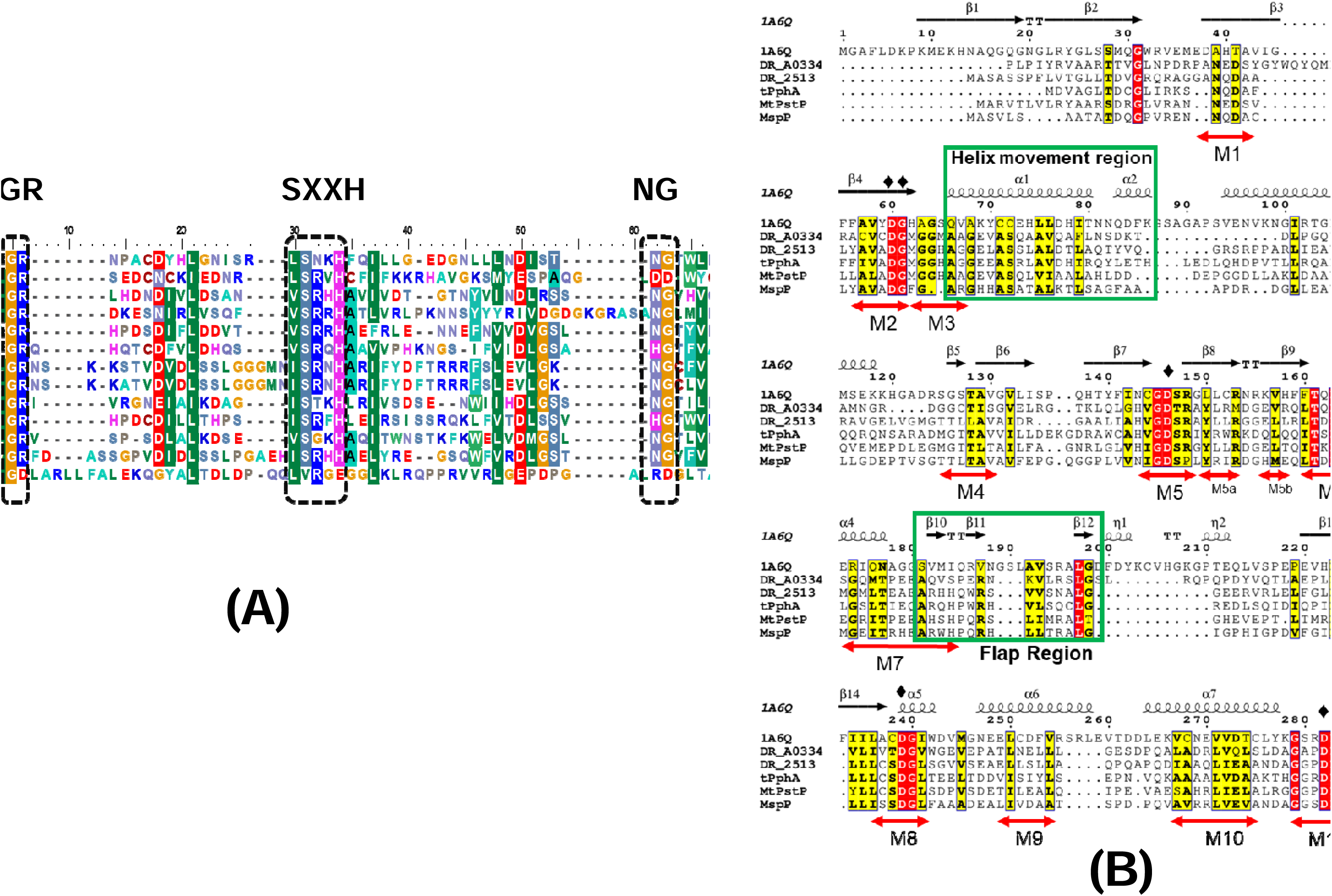
Comparative sequence analysis of FHA domain and PP2C-type Ser/Thr phosphatases. (A) Multiple sequence alignment (MSA) analysis of FHA domain proteins and showing conservation along conserved motifs (GR, SXXH, NG). (B) MSA of *D. radiodurans* PP2C-type phosphatases (DR2513 and DRA0334) with representative bacterial and eukaryotic homologs highlighting the conserved PP2C signature motifs (M1–M11).

STRING analysis of DRA0334 and DR2513 STPPs protein suggests their role in dephosphorylation of key STPKs of *D. radiodurans* (Fig. S3). The interaction network reveals strong associations with several STPKs, including DRA0332 (score: 0.927), DR0058 (0.872), DR0534 (0.867), and DR1243 (0.857), implying that DRA0334 may function in concert with these kinases to modulate phosphorylation states within the cell. Notably, its association with RqkA (DR2518) (score: 0.860), a kinase involved in radiation resistance and DNA double-strand break repair, suggests DRA0334 STPP role in the *D. radiodurans* stress response mechanisms. Interestingly, STRING analysis also predicted interaction with operon partner FraH-related protein (DRA0333) and a STPK (DRA0332) strengthens the likelihood of a shared regulatory or functional pathway (as detailed in subsequent results sections).

Another predicted STPP, DR2513 STRING interaction network indicates strong associations between DR2513 STPP and several STPKs, including DR1243, DR0058, and DR1213, with confidence scores ranging from 0.877 to 0.900 (Fig. S6). Although STRING interaction data are useful for predicting functional associations between STPPs (DRA0334 and DR2513) and STPKs in *D. radiodurans*, experimental validation of both functional and physical interactions is essential to strengthen these predictions and confirm the proposed dephosphorylation-dependent regulatory networks. Therefore, the predicted STPP–STPK interactions were further examined through subsequent in vivo and in vitro experiments and discussed in results 3 section.

The phosphatases belonging to the PPM family, act on serine/threonine residues and require Mg^2+^ or Mn^2+^ metal ions. Bacterial PPM family phosphatases share a common catalytic domain with the eukaryotic PP2C. Among bacterial PPMs, two subfamilies: One subfamily, lacks motifs 5a and 5b within the catalytic domain and includes phosphatases linked to non-eukaryotic serine/threonine kinases exemplified by *B. subtilis* sporulation-specific phosphatase SpoIIE and stress response phosphatases RsbU, RsbP and RsbX [1]. The second subfamily, denoted as eukaryotic type STPPs, encompasses members containing all 11 signature including 5a and 5b motifs and represents cognate phosphatases to eukaryotic serine/threonine kinases (eSTKs) exemplified by PrpA (*Salmonella enterica* serovar) [49], PrpC (*B. subtilis*)[7], PrpC (*Mycoplasma pneumoniae*)[50], PstP (*M. tuberculosis*) [8], tPphA (*Thermosynechococcus elongates*) [51], Pph1 (*Myxococcus xanthus*) [52], Stp1 (*Staphylococcus aureus*) [53], PhpP (*Streptococcus pneumoniae*) [54] (Fig. 1B).

The Multiple sequence alignment (MSA) of DR2513 and DRA0334 proteins with the human (PDB code 1A6Q) and other bacterial PP2C phosphatase [55] reveals that both DR2513 and DRA0334 proteins possess 11 characteristic motifs of the PP2C family, including motifs 5a and 5b [56] (Fig. 1B). Both *D. radiodurans* specific PP2C phosphatases (DR2513 and DRA0334) have methionine (M) instead of histidine (H) at position 62 of human PP2C phosphatase. This position is known to play a role in breaking the P-O bond. Similar type of replacement of amino acid is also found in PstP of *M. tuberculosis* and PphA of *T. elongatus* (Fig. 1B) suggesting the functional conservation this residues among bacterial STPPs. For PPM type phosphatases, motifs (M) 1, 2, 5, and 11 are essential for catalytic phosphatase activity, being involved in metal ion coordination. Additionally, helix α1 and 2 are the part of helix movement region crucial for the substrate specificity while the flap region remain mobile and involve in substrate binding and catalytic activity [57] (Fig. S4). Homology models of the catalytic domains support these findings and showed that model of DR2513 and DRA0334 show typical bacterial structural similarities with *M. tuberculosis* PstP having conservation in residue for the coordination of a metal ion, residues involve in splitting the P-O bond, and the repositioning of the flap region [58] (Fig. S4).

### 2. Biophysical and Catalytic parameter analysis of PP2C type protein phosphatase (DRA0334)

The wild-type DRA0334 and two of its point mutants for the prospective residues involve metal coordinating aspartate (D394A and D419A) were purified to its homogeneity (Fig. S5). The CD spectra showed that wild-type DRA0334 and its mutants have defined secondary structures and replacement of key catalytic aspartic acid residue D394 and D419 with alanine did not affect the secondary structures folding (Fig. S6). Biochemical characterization confirmed that DRA0334 is a functional protein phosphatase (Fig. 2). Using the synthetic substrate p-nitrophenyl phosphate (pNPP), purified DRA0334 showed a concentration-dependent increase in phosphatase activity, demonstrating its enzymatic function (Fig. 2A). DRA0334 mutants (D394A and D419A) showed no hydrolysis of pNPP, highlighting the essential role of this residue for catalytic activity (Fig. 2B). DRA0334 was found to be a Mn^2+^-dependent phosphatase but also showed phosphatase activity with Co^2+^. No phosphatase activity was observed in the presence of Zn^2+^, Cu^2+^, Ni^2+^, Ca^2+^, and Mg^2+^ using the pNPP assay (Fig. 2C), a finding consistent with previously characterized bacterial PP2Cs such as PphC [59] and PstP [60]. Optimal activity with pNPP was observed at 4-6 mM MnCl_2_ (Fig. 2D). The chelator EDTA caused a significant decrease in DRA0334 activity, further supporting its dependence on a bivalent metal cation. Kinetic analysis using pNPP showed that DRA0334 has observed Km and Vmax values are 3.27 mM and 507.09 pmol min^-1^ µg^-1^ respectively while Kcat (turnover number) 0.5507 s^-1^and Kcat/Km (catalytic efficiency) is 0.168 s^-1^ mM^-1^ (Fig. E, F).

**Figure 2.**
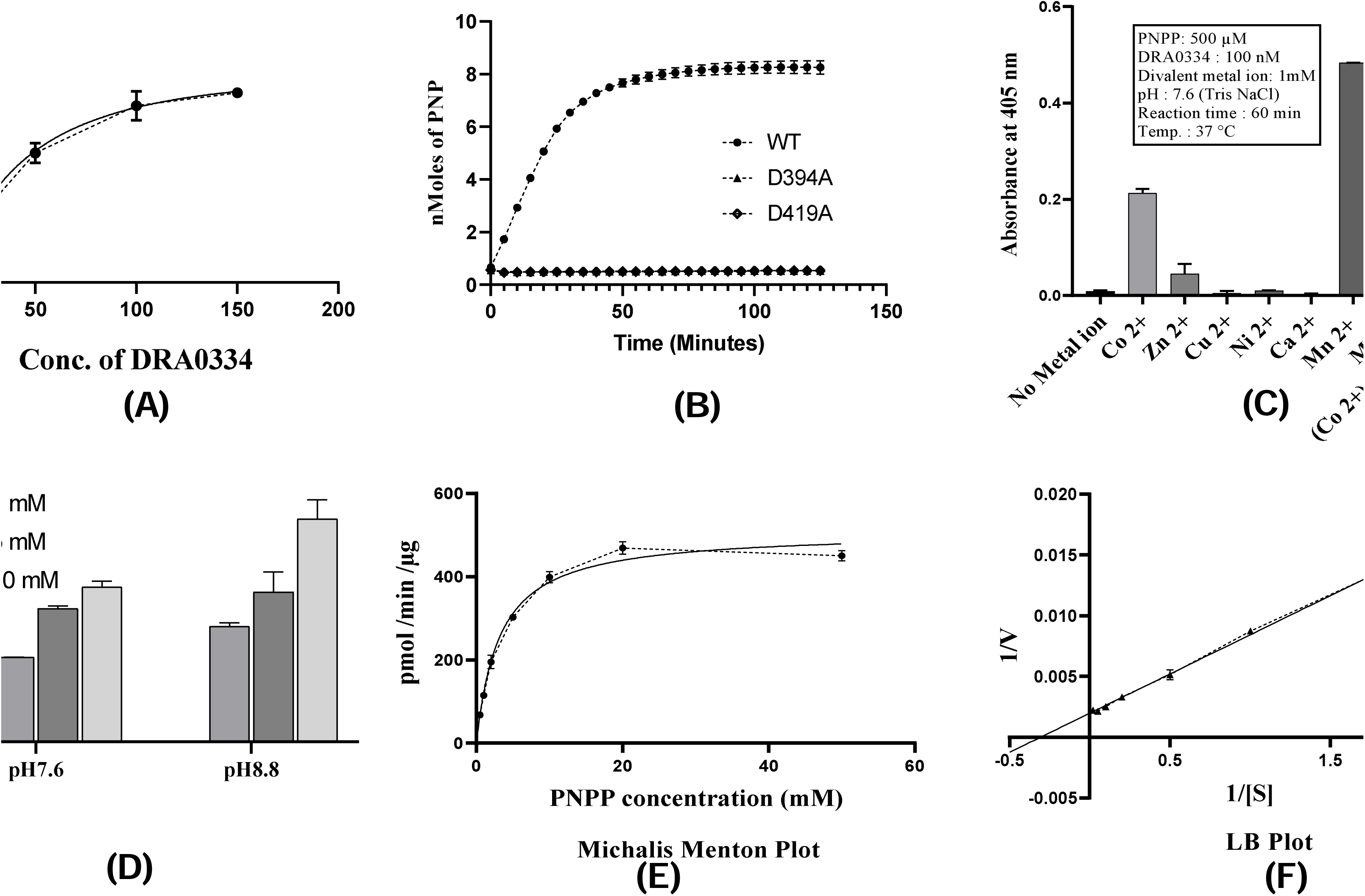
Biochemical characterization of DRA0334 phosphatase activity. (A) Concentration-dependent phosphatase activity of DRA0334 using pNPP as substrate. (B) Time-dependent pNPP hydrolysis by wild-type DRA0334 (WT) and its catalytic mutants (D394A and D419A). (C) Metal ion dependency showing activity with Mn²□ and Co²□. (D) Optimal MnCl□ concentration for activity. (E–F) Kinetic parameters (Km, Vmax, Kcat) derived from Michaelis–Menten and LB plot analysis. Data are presented as the mean ± standard error (SE) from three independent experiments for the each panel.

### 3. STPPs (DRA0334 and DR2513) may have broad specificity for *D. radiodurans* STPKs

STPPs typically exhibit broad substrate specificity toward their cognate serine/threonine protein kinases (STPKs), thus playing essential roles in regulating reversible phosphorylation in bacteria. For instance, PrpC STPP of *B. subtilis* dephosphorylates the STPK PrkC [61]; PstP STPP of *M. tuberculosis* targets several STPKs, including PknA [8], PknB [62], PknH [8], and PknJ [63]; STPP like PhpP in *S. pneumoniae* dephosphorylates StkP [64]; and Stp1 in *Staphylococcus aureus* acts on StkP [9]. These examples collectively highlight the widespread functional interplay between bacterial STPKs and STPPs in maintaining the dynamic balance of phosphorylation-mediated signaling networks.

STRING analysis of DRA0334 and DR2513 STPPs shows they may play a role in dephosphorylating key STPKs in *D. radiodurans*. The network reveals strong interactions with several kinases, such as DRA0332, DR0058, DR0534, and DR1243, indicating a coordinated regulation of cellular phosphorylation (Fig. S3). Importantly, the interaction with RqkA (DR2518) connects DRA0334 to radiation stress response and DNA repair pathways. Predicted links with FraH-related protein (DRA0333) and DRA0332 further support a shared functional or regulatory module (Fig. S3). Therefore, taking lead from STRING analysis both PP2C-type STPPs (DR2513 and DRA0334) ability to dephosphorylate specific STPKs was investigated in vivo and in vitro. For in vivo dephosphorylation assays, co-expression of STPKs (RqkA and DR1243) with STPPs (DRA0334 and DR2513) was carried out in *E. coli* surrogate hosts BL21 and Novablue using inducible (pET28a) and constitutive (pKNT) expression systems, respectively. The level of dephosphorylation of STPKs by these STPPs was monitored by detecting changes in the intensity of phosphorylated protein bands using an anti-Ser/Thr antibody (Fig. 3A). In *E. coli* BL21 cells, DR2513 dephosphorylated approximately 50% of phosphorylated RqkA and DR1243 STPKs, whereas DRA0334 exhibited stronger activity, reducing phosphorylation levels of RqkA by nearly 60% and DR1243 by 80% (Fig. 3A). Notably, the in vivo phosphatase activity was even higher in the Novablue strain, where STPPs were expressed from the constitutive pKNT vector while cognate RqkA STPK expressed from constitutive pRAD vector. Under these conditions, DRA0334 dephosphorylated ∼80% and DR2513 ∼60% of RqkA, respectively (Fig. 3B). This difference in dephosphorylation efficiency between the two *E. coli* hosts likely reflects variations in STPKs and STPPs expression kinetics, redox status, or metabolic conditions influenced by the nature of the expression system employed like inducible (pET28a) versus constitutive (pKNT). Notably, under both experimental conditions, DRA0334 exhibited comparatively higher dephosphorylation activity than DR2513. Although the precise reason for this difference remains unclear, a plausible explanation is that DRA0334 possesses an N-terminal KI-FHA domain (Fig. S2 B), which is known to recognize phosphothreonine residue. This domain may enhance substrate specificity and binding affinity toward phosphorylated STPKs, thereby facilitating more efficient dephosphorylation than DR2513. However, this hypothesis requires further validation through dedicated experiments, which are described in subsequent results sections. Collectively, these findings indicate that both DRA0334 and DR2513 specifically dephosphorylate *D. radiodurans* STPKs (RqkA and DR1243), with DRA0334 potentially exerting more targeted regulatory control due to its modular KI-FHA architecture.

**Figure 3.**
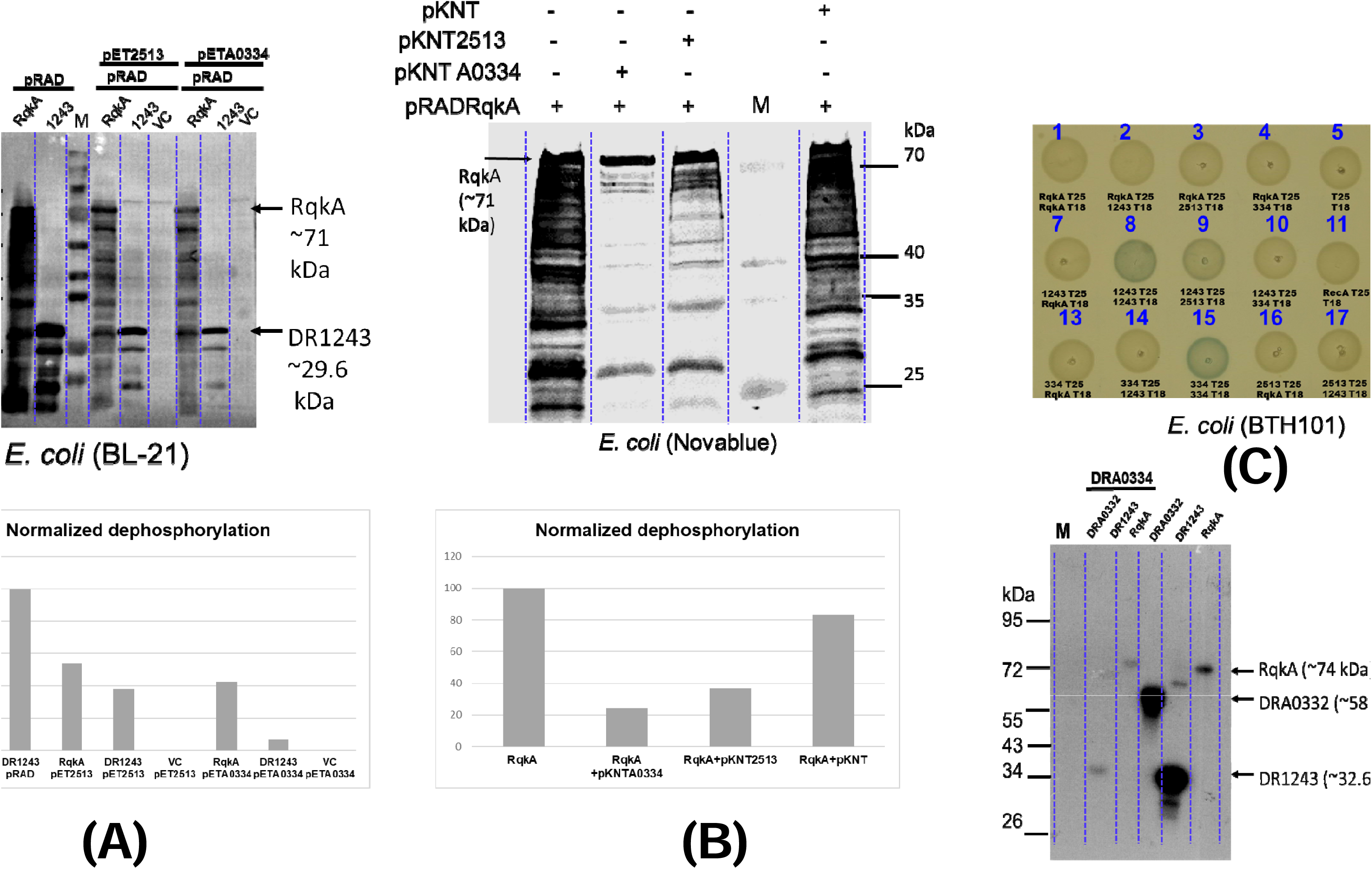
Functional interaction and dephosphorylation activity of STPPs (DRA0334 and DR2513). (A & B) In vivo and Ex vivo dephosphorylation of RqkA and DR1243 STPKs. Blot showing dephosphorylation of RqkA, RecA, and DR1242 STPKs by DRA0334 and DR2513 STPPs. (C) Protein-protein interaction of STPKs-STPP demonstration in *E. coli* BTH strain. (D) DRA0334 catalyzed dephosphorylation of three STPKs (RqkA, DRA0332, and DR1243) shown in vitro using [^32^P] labelled STPKs. The data shown are representative of three independent experiments for each panel. For panels A and B, densitometric analysis of Western blot signals was performed on a single representative blot using the ImageJ software. Signal intensities were quantified after background subtraction and used for normalization.

Interestingly, bacterial two-hybrid assays revealed no stable interactions between the STPPs and their target kinases, suggesting that these interactions may be transient and catalytic in nature (Fig. 3C). However, self-interaction of STPPs was evident, particularly for DRA0334 (spot no. 15) and DR2513 (spot no. 18) (Fig. 3C), indicating possible oligomerization that may influence their activity or stability.

To further validate the Fig. 3A, B in vivo results, in vitro phosphatase assays were conducted using purified recombinant proteins. Two STPPs (DRA0334 and DR2513) and three STPKs (RqkA, DRA0332, and DR1243) of *D. radiodurans* were expressed and purified from *E. coli* BL21 (Fig. S5). Unfortunately, DR2513 accumulated largely as an inclusion body, and refolded DR2513 failed to show phosphatase activity in vitro. In contrast, DRA0334 was soluble and exhibited Mn^2+^-dependent phosphatase activity, displaying kinetic parameters on synthetic substrate comparable to other bacterial PP2C-type phosphatases (Fig. 2). Remarkably, DRA0334 efficiently dephosphorylated all three STPKs at a ∼1:3 STPP:STPK molar ratio (Fig. 3D), reinforcing its broad substrate specificity within the *D. radiodurans* kinase network. Altogether, these findings provide strong evidence that *D. radiodurans* STPPs (DRA0334 and DR2513) play key roles in reversing the phosphorylation state of STPKs, thereby may contributing to the dynamic control of signaling pathways of *D. radiodurans*. The reversible phosphorylation mechanism managed by these kinases particularly RqkA and phosphatases is likely crucial for *D. radiodurans* capacity to coordinate DNA repair and cell division processes after radiation-related damage and need separate investigation as function of RqkA in DNA damage response and cell cycle control is well documented [26, 34, 36, 42, 65].

### 4. FHA / KI-FHA docking interactions in modulating STPK dephosphorylation dynamics

FHA domains are known to enhance kinase activity by functioning as phospho-dependent docking and regulatory modules. Mechanistically, FHA domains recognize specific phosphothreonine (pThr)-containing motifs on adaptor or substrate proteins, thereby facilitating the assembly of productive kinase–substrate complexes [16]. The KI-FHA domain is a specialized variant of the FHA domain that binds phosphorylated threonine residues and is involved in signaling networks regulating cell growth and division, as exemplified by the KAPP phosphatase in plants and the Ki-67 protein in humans [21, 66]. In *D. radiodurans*, DRA0333 harbors a well-defined FHA domain, whereas DRA0334, a PP2C-type phosphatase, contains a KI-FHA domain (residues 210–368) located between an N-terminal unstructured region (residues 1–124) and its PP2C catalytic domain (residues 369–614) (Fig. S2 B). To investigate the functional roles of these domains, DRA0333 (FHA domain) and DRA0334 (KI-FHA PP2C phosphatase) were tested for their effects on the phosphorylation and dephosphorylation of *D. radiodurans* serine/threonine protein kinases (STPKs: RqkA, DR1243, and DRA0332) and myelin basic protein (MBP) (Fig. 4).

**Figure 4.**
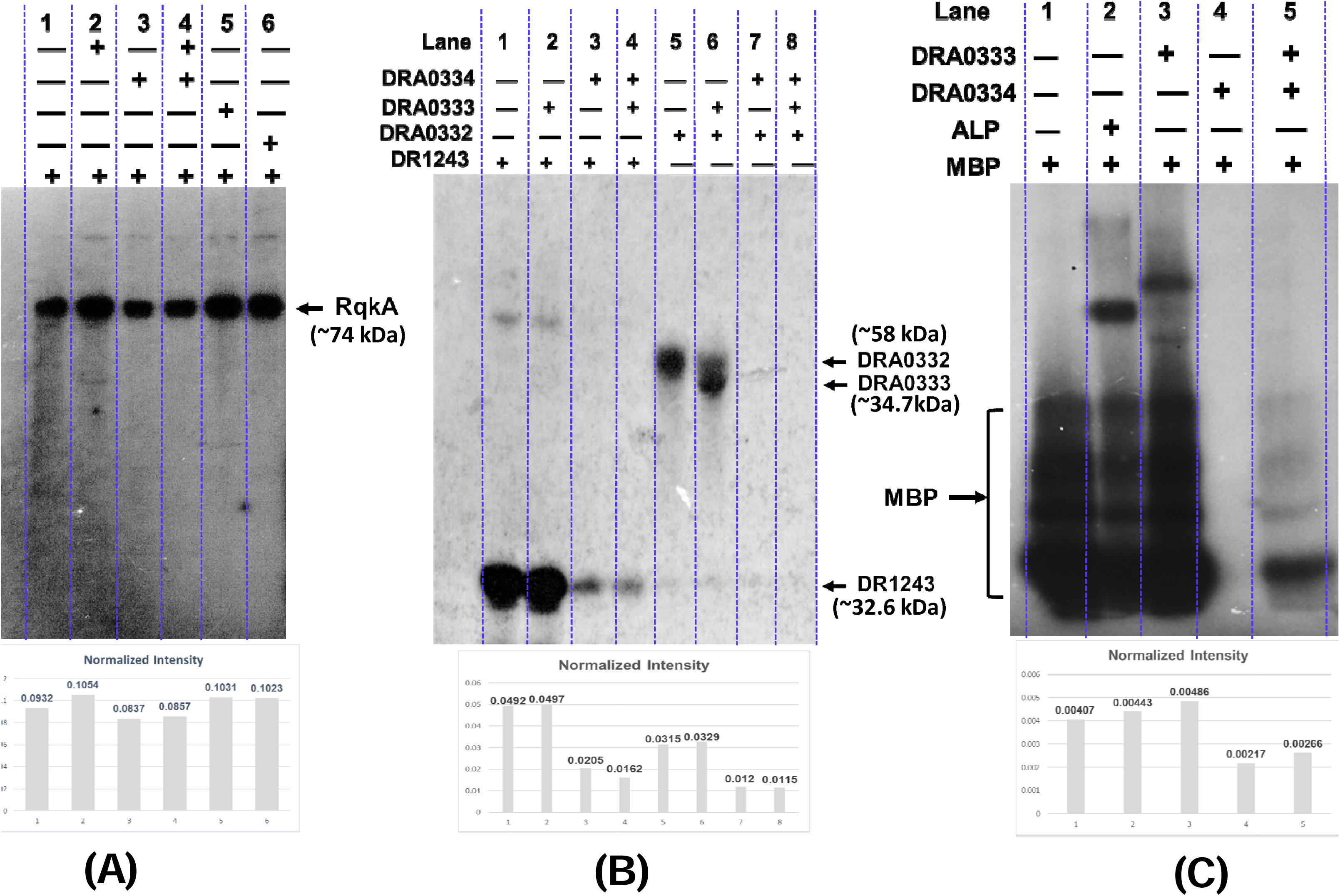
Functional interplay between DRA0333 (FHA protein) and DRA0334 (PP2C phosphatase). (A) Dephosphorylation of RqkA STPK by DRA0334 phosphatase or by its mutants (D394A and D419A) in the presence of operonic partner FHA domain protein DRA0333. (B) Dephosphorylation of DRA0332 and DR1243 STPKs by DRA0334 phosphatase in the presence of operonic partner FHA domain protein DRA0333. (C) Dephosphorylation of MBP phosphoprotein by DRA0334 phosphatase in the presence of operonic partner FHA domain protein DRA0333. Data are presented as representative autoradiograph of three independent experiments for the each panel. For panels A to C, densitometric analysis of autoradiographs signals was performed on a single representative autoradiograph using the ImageJ software. Signal intensities were quantified after background subtraction and used for normalization.

A good increase in the autophosphorylation of RqkA was observed in the presence of FHA domain protein DRA0333 (Fig. 4A, lane 2), consistent with the canonical stimulatory role of FHA domains. DRA0333 also produced a modest enhancement in DR1243 phosphorylation (Fig. 4B, lane 2). Interestingly, DRA0332 appeared to phosphorylate DRA0333 itself, as indicated by the reduction in DRA0332 signal and a corresponding increase in DRA0333 phosphorylation (Fig. 4B, lane 6), suggesting a phosphotransferase-type interaction rather than kinase activation. When RqkA-mediated MBP phosphorylation was examined, DRA0333 markedly enhanced transphosphorylation efficiency (Fig. 4C, compare lanes 1 and 3), supporting the role of FHA domains as scaffolds that facilitate oligomerization and signal amplification through phospho-dependent binding. Collectively, these results confirm the classical function of FHA domain proteins as positive regulators of kinase signaling by promoting phosphorylation events.

In contrast, the PP2C phosphatase DRA0334 effectively dephosphorylated RqkA, DR1243, and DRA0332 with varying efficiency (Fig. 4A, lane 3; Fig. 4B, lanes 3 and 7, and Fig. 3D). However, the KI-FHA domain of DRA0334 displayed a unique regulatory feature. Dephosphorylation of RqkA and phosphorylated MBP (generated by incubation with RqkA kinase as described in Methods) by DRA0334 was markedly inhibited in the presence of DRA0333 (Fig. 4A, lane 4; Fig. 4C, lane 5), whereas dephosphorylation of DR1243 and DRA0332 remained unaffected (Fig. 4B, lanes 4 and 8). These observations suggest that perhaps the FHA domain of DRA0333 competes with the KI-FHA domain of DRA0334 or vice versa for binding to phosphorylated residues on RqkA or MBP, thereby stabilizing RqkA in its active conformation and promoting productive auto- and trans-phosphorylation. Furthermore, such KI-FHA–mediated interactions may also enhance substrate access to the catalytic phosphatase site of DRA0334. Consistent with this interpretation, DRA0334 displayed higher in vivo dephosphorylation efficiency toward RqkA compared to DR2513 (see Fig. 3A & B), likely due to its KI-FHA-mediated phosphosite recognition. Thus, the FHA domain of DRA0333 likely competes with the KI-FHA domain of DRA0334 for substrate docking, thereby weakening KI-FHA-mediated stimulation of RqkA autophosphorylation. This competitive interaction also inhibits the KI-FHA–associated PP2C phosphatase activity of DRA0334. Nonetheless, this represents a novel and previously unexplored observation in which both kinases and phosphatases employ FHA domain–mediated docking to recognize phosphosites and modulate their enzymatic activities.

Further evidence supporting the functional importance of the KI-FHA domain comes from analysis of DRA0334 catalytic mutants (D394A and D419A), which are defective in dephosphorylation (Fig. 2B). These mutants instead enhanced RqkA autophosphorylation relative to wild-type DRA0334 (Fig. 4A, lanes 5–6 vs. 3). This suggests that, in the absence of phosphatase catalytic activity, the KI-FHA domain can mimic an FHA-like stimulatory effect by stabilizing kinase–substrate interactions. As a control, alkaline phosphatase (ALP) was included as a standard phosphatase and dephosphorylated MBP (Fig. 4B, lane 2). Interestingly, residual activity of RqkA (used for the generation of phosphorylated MBP) was able to phosphorylate ALP itself, producing a strong autoradiographic signal at the corresponding molecular weight (Fig. 4C, lane 2, ALP arrow).

Taken together, these findings demonstrate that the FHA / KI-FHA domain enhances both the auto- and transphosphorylation activity of *D. radiodurans* RqkA STPKs, while DRA0334’s phosphatase activity depends on its PP2C catalytic site integrity and is modulated by competitive interaction between its KI-FHA domain and FHA domain of DRA0333. This finding reveals a previously unrecognized regulatory mechanism wherein both kinases and phosphatases exploit FHA / KI-FHA domain–mediated phosphosite docking to fine-tune their catalytic activities. Notably, of the three STPKs examined, only RqkA exhibited this novel mode of regulation. This highlights a STPK-specific regulatory interplay between DRA0333 and DRA0334, wherein competition between FHA / KI-FHA domains fine-tunes the kinase–phosphatase balance. Such a mechanism provides new insight into the dynamic control of STPK-mediated signaling in *D. radiodurans*.

### 5. The DRA0334 modular variants confirms the regulatory crosstalk of FHA / KI-FHA domains in kinase–phosphatase antagonism

The preceding section data shows that the interaction between DRA0333 and DRA0334 represents a novel regulatory mechanism, where FHA / KI-FHA domain competition balances the activities of kinases and phosphatases as both kinases and phosphatases uses these domains-mediated phosphosite binding to adjust their catalytic activities in dynamic fashion. However, the significance of divergent FHA domain (KI-FHA) fusion with PP2C phosphatase domain of DRA0334 is not clear about why divergent KI-FHA domain exist with PP2C instead true FHA domain as found in DRA0333 operon partner protein. Therefore, it was envisaged that domain swapping of KI-FHA with FHA domain in DRA0334 might impact the functional properties DRA0334 phosphatase in more radical way instead as observed for KI-FHA in results shown in figure 4. Therefore to strengthen the hypothesis of how the FHA / KI-FHA module affects PP2C activity, we created three DRA0334 constructs and looked at their ability to dephosphorylate two phosphoprotein substrates: the STPK RqkA and RecA (Fig. 5A). The constructs included: (i) full-length DRA0334 (wild-type) with native KI-FHA and PP2C domains; (ii) PP2C-only, which had only the catalytic PP2C domain; and (iii) a chimera, where the native KI-FHA was replaced by the FHA domain of DRA0333 fused to the PP2C catalytic domain (Fig. S7).

**Figure 5.**
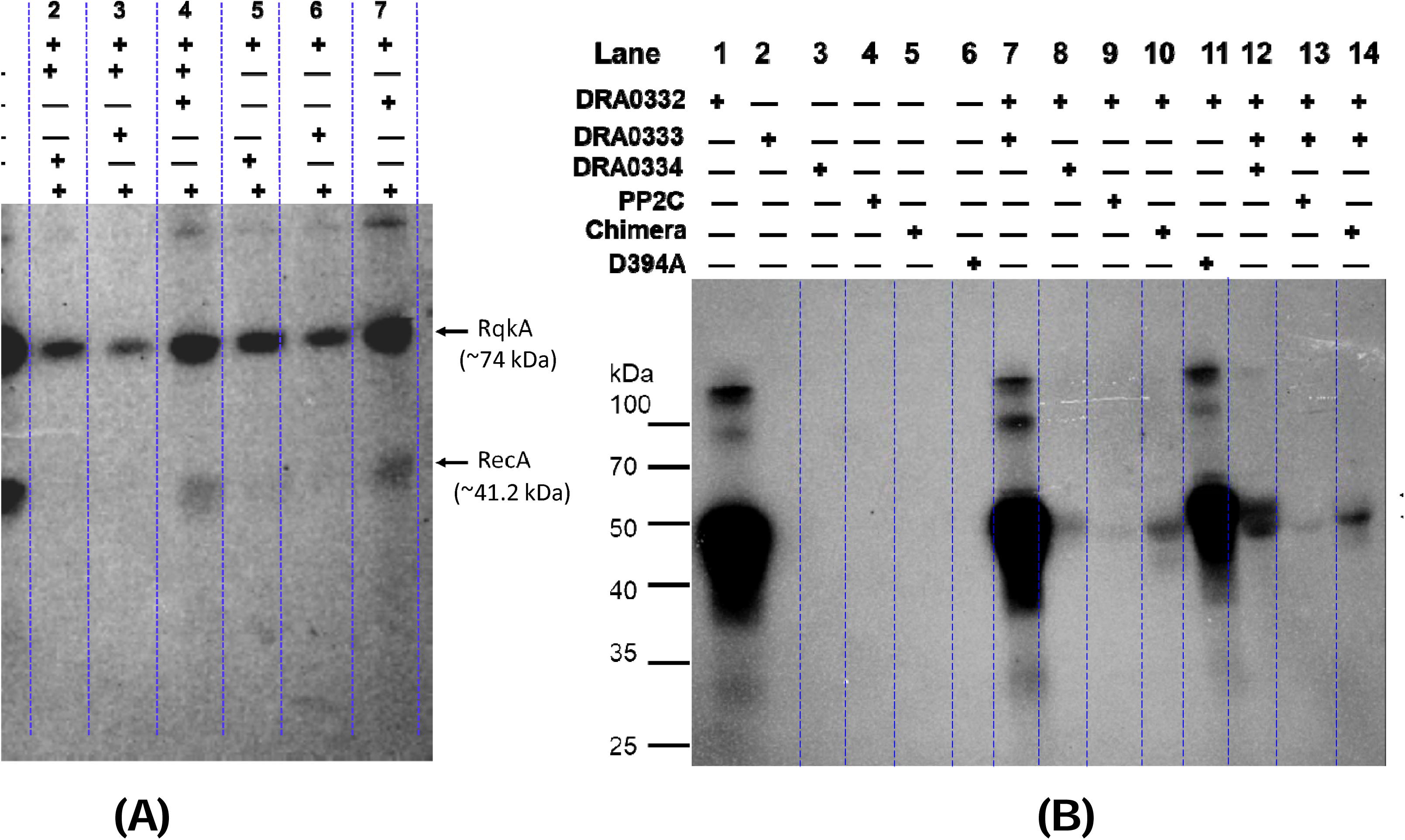
Regulatory role of KI-FHA domain of DRA0334 PP2C phosphatase in dephosphorylation of phosphoproteins. The role of the N-terminal KI-FHA module of DRA0334 in modulating phosphatase activity was examined using STPKs; RqkA and DRA0332 and the phosphoprotein RecA as substrates. Three DRA0334 constructs were employed: (i) full-length wild-type DRA0334 containing the native KI-FHA and PP2C domains; (ii) a PP2C-only construct comprising the catalytic PP2C domain; and (iii) a chimeric construct in which the native KI-FHA module was replaced with the FHA domain of DRA0333 fused to the PP2C catalytic domain. The ability of these constructs to dephosphorylate the phosphoprotein substrates was assessed for (A) RqkA and RecA and (B) DRA0332. Dephosphorylation was analyzed by autoradiography, and the data shown are representative autoradiographs from three independent experiments for each panel.

These variants were employed for the dephosphorylation of phosphorylated substrate proteins (RqkA and RecA) and analyzed the reactions using autoradiography (Fig. 5A). Data shows that as expected the wild-type DRA0334 showed modest dephosphorylation of RqkA kinase which is modestly improved by addition of DRA0333 FHA domain corroborating results reported in Fig. 4A (see RqkA, lane 4 & 7 compare to control lane 1, Fig. 5A). This result can also be interpret in the way that the KI-FHA domain of DRA0334 exerts a stimulatory effect on RqkA autophosphorylation while simultaneously suppressing the PP2C phosphatase activity of DRA0334 (RqkA, lane 7 vs. control lane 1; Fig. 5A). In contrast, the presence of the FHA domain of DRA0333 appears to compete with the KI-FHA domain for RqkA interaction, thereby reducing kinase stimulation and allowing the PP2C phosphatase activity of DRA0334 to dominate, resulting in enhanced dephosphorylation of RqkA (RqkA, lane 4 vs. lane 7; Fig. 5A). A similar trend was observed for RecA, where DRA0334-mediated dephosphorylation was modestly enhanced in the presence of DRA0333 (RecA, lanes 4 and 7 vs. lane 1; Fig. 5A). Interestingly, the most efficient dephosphorylation of both RqkA and RecA, reflected by the lowest autoradiographic signal, was observed with the PP2C-only construct (lane 6, Fig. 5A), whereas the presence of the DRA0333 FHA domain did not affect dephosphorylation (lane 3, Fig. 5A). Collectively, these results indicate that the KI-FHA domain, either alone or in association with an FHA-domain protein, can independently compete for binding to phosphorylated residues on RqkA and RecA. This FHA / KI-FHA–mediated phosphosite docking correlates with increased phosphorylation of RqkA/RecA while simultaneously restricting productive catalysis by the PP2C phosphatase domain. In contrast, in the absence of FHA / KI-FHA domains, the PP2C domain of DRA0334 dephosphorylates its substrates efficiently without steric or competitive interference.

Additionally, although the isolated PP2C phosphatase domain achieved complete or near-complete dephosphorylation, the chimeric construct exhibited higher dephosphorylation efficiency than the wild-type but lower than the PP2C-only construct, as indicated by an intermediate autoradiographic signal (Lane 5, Fig. 5A). Notably, the chimera comprising the PP2C domain fused to the FHA domain of DRA0333 showed nearly comparable dephosphorylation of both RqkA and RecA when supplemented with DRA0333, rather than stimulating RqkA phosphorylation (Lanes 2 and 5, Fig. 5A). These observations suggest that the native KI-FHA domain of DRA0334 confers specificity toward RqkA phospho-docking sites compare to FHA domain of DRA0333, and highlight dynamic processing of RqkA by two opposing activities embedded within a single DRA0334 polypeptide: the KI-FHA domain, which facilitates RqkA phosphorylation through substrate docking, and the PP2C domain, which catalyses RqkA dephosphorylation (compare Lanes 6 and 7, Fig. 5A). Thus, the phosphatase catalytic domain alone is insufficient for optimal and dynamic substrate processing in the absence of a docking module, supporting a push–pull regulatory mechanism of phosphorylation and dephosphorylation.

These results were further verified on operon partner STPK (DRA0332) (Fig. 5B). Here, again PP2C-only showed near complete dephosphorylation of DRA0332 STPK, followed by wild-type DRA0334 and chimera (lane 8, 9, and 10, Fig. 5B). The dephosphorylation of DRA0332 STPK was largely unaffected for PP2C-only and chimera by addition of DRA0333 FHA domain protein while wild-type DRA0334 activity was inhibited (lane 12, 13, and 14, Fig. 5B). A phosphorylated band of DRA0333 also visible in these lanes due to phosphotransferase activity by DRA0332 STPK (as also observed in lane 6, Fig. 4B). Additionally, various reaction control taken in lane 1 to 6, Fig. 5B. Interestingly, the phosphorylation-stimulatory effect of the KI-FHA domain on RqkA observed in Fig. 5A was not evident for the operonic STPK DRA0332. Instead, the FHA domain of DRA0333 elicited a comparable stimulatory effect on DRA0332 (compare lanes 8 and 12 in Fig. 5B), indicating STPK-specific interactions of the KI-FHA and FHA domains.

Overall, these results show that efficient dephosphorylation by DRA0334 needs both docking module (KI-FHA) and PP2C catalytic domain. Losing the docking module (PP2C-only) or replacing it with a different FHA (chimera) may change substrate access / or interaction / or positioning, leading to loss of fine-tuned dephosphorylation of RqkA, DRA0332 STPKs or substrate phosphoprotein (RecA). This pattern aligns with a model where the KI-FHA or FHA domain provides STPK specificity and form a dynamic regulatory circuit for the kinase-phosphatase interaction and resetting of signaling (Fig. 6).

**Figure 6.**
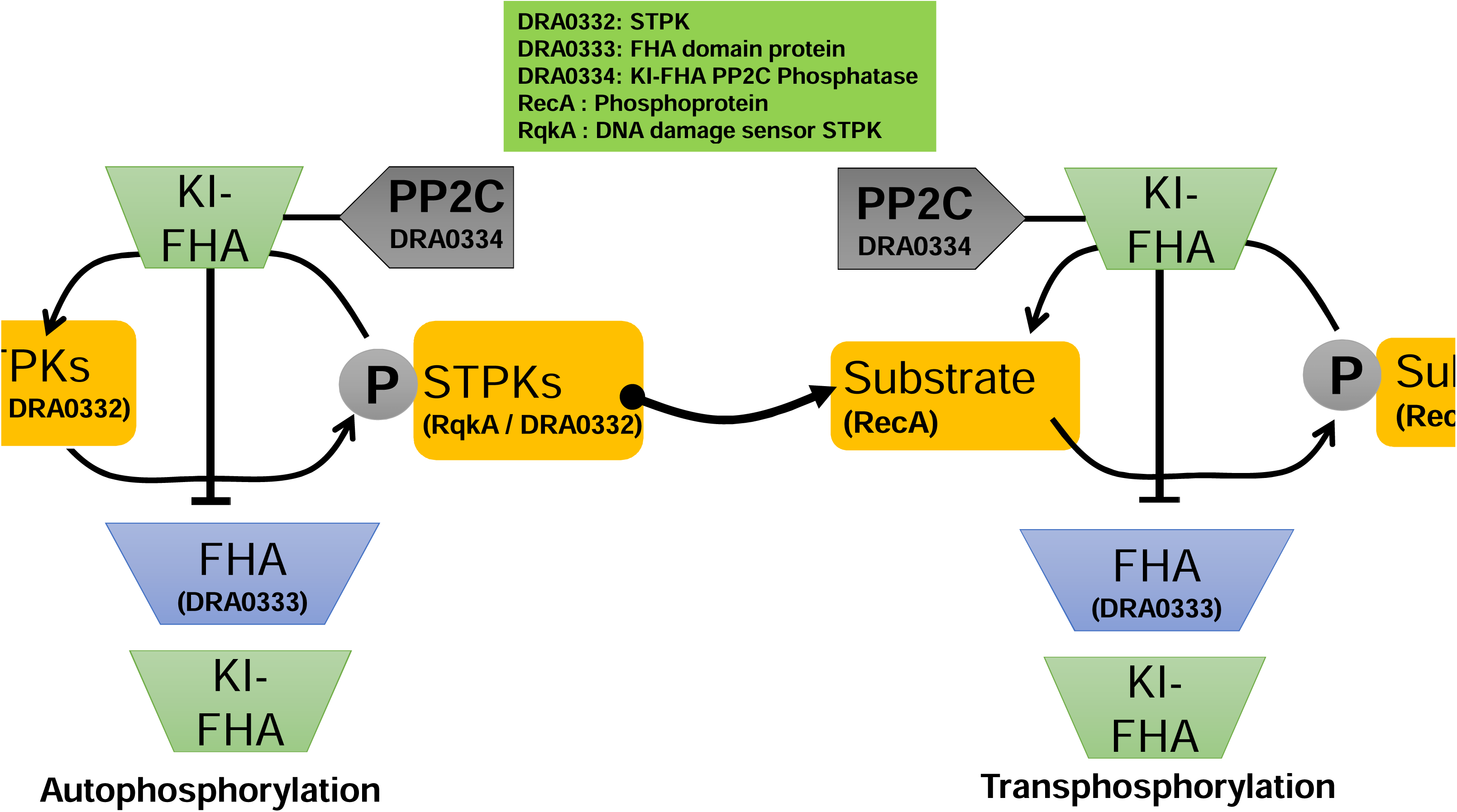
Proposed model for KI-FHA and FHA domain interplay in dynamic regulation of kinase and phosphatase activity. The model summarizes a dynamic regulatory circuit where the FHA-domain protein (DRA0333) or alone KI-FHA module enhances the phosphorylation cycle by acting as a phospho-dependent docking and regulatory module, stabilizing active kinase conformations and promoting autophosphorylation of STPKs (RqkA / DRA0332) or trans-autophosphorylation of cognate substrate (RecA). In contrast, KI-FHA domain when fused with the PP2C-type phosphatase (DRA0334) counterbalances this effect through its possible competitive titration of FHA domain proteins. These features of KI-FHA fused PP2C phosphatase (DRA0334) may enables phospho-substrate recognition and precise dephosphorylation of kinases or phosphoproteins. This dual-domain configuration allows DRA0334 to act as both a catalytic phosphatase and a phospho-signal modulator. The model depicts how competitive binding between operonic partner protein; the FHA domain of DRA0333 and the KI-FHA domain of DRA0334 regulates kinase–phosphatase equilibrium, thereby employing a novel way of fine-tuning phosphorylation signaling. Such interplay ensures transient and reversible control of STPK-STPP mediated signaling events, crucial for coordinating processes like DNA repair, cell division, and stress recovery. The inset shows the structural domains involved like canonical FHA domain, KI-FHA domain, and PP2C catalytic core.

## Discussion

FHA domains are conserved phosphothreonine-binding modules that mediate phospho-dependent protein–protein interactions essential for signaling and regulation. They are highly enriched in DNA damage response and genome maintenance proteins, where they recognize damage-induced phosphothreonine motifs to promote kinase activation, autophosphorylation, and assembly of repair complexes. Structurally, FHA–pThr interactions generate orientation-specific interfaces that increase local kinase–substrate concentration and stabilize catalytically active conformations. FHA domains can also regulate intramolecular autoinhibition, thereby switching kinases from inactive to active states [67]. Acting as modular scaffolds, they enhance signal amplification and pathway specificity, with binding preferences around pT±1 to 3 residues dictating distinct downstream outcomes (Fig. S1 A) [68]. FHA–STPK interactions are widely reported in bacteria, underscoring their central role in controlling kinase activity and substrate phosphorylation [46, 69, 70].

In this study, a PP2C-type phosphatase DRA0334 is identified as the specific phosphatase for the DNA damage sensor kinase RqkA and other STPKs of *D. radiodurans* (Fig. 3). DRA0334 displays the classic metal ion-dependent catalytic features common to bacterial PP2C family members (Fig. 2), but it also has a unique N-terminal KI-FHA domain that helps recognize substrates and regulate interactions (Fig. 4, 5, & S2). Data indicate that DRA0334 dephosphorylates STPKs like RqkA, DR1243, and DRA0332 in a Mn²□-dependent way (Fig. 3). However, its activity is influenced by the neighboring operon partner DRA0333 (a FHA domain protein) (Fig. 4). The contrast between the phosphorylation stimulation by DRA0333 and the dephosphorylation by DRA0334 creates an operon-driven circuit that uniquely exist in the *D. radiodurans* where genes encoding FHA domain protein (DRA0333), STPK (DRA0332), and KI-FHA–PP2C phosphatase (DRA0334) are clustered represents a unique and previously unreported arrangement suggestive of a novel regulatory module (Fig. S2). This setup may allow for rapid adjustments in phosphorylation levels and ensures localized regulation through modular domain interactions instead of long-range signaling intermediates. Additionally, the presence of a von Willebrand A (vWA) domain protein, DRA0331, within the same operon hints at a possible scaffolding or localization role. This could connect the kinase-phosphatase-FHA module to DNA repair sites or membrane areas, consistent with the modular design seen in hyperthermophilic archaeon [46].

The domain architecture of DRA0334 closely resembles that of KAPP-like PP2C phosphatases in *A. thaliana*, which regulate receptor-like kinases (RLKs) such as SERK1, CLV1, and FLS2, thereby controlling signaling pathways involved in meristem development and receptor-mediated phosphorylation events [21, 71, 72]. This structural and functional analogy suggests that DRA0334 and its operonic partners may constitute an evolutionarily adapted bacterial signaling circuit analogous to plant FHA–PP2C–kinase regulatory systems. The KI-FHA domain of DRA0334 seems to has a dual function. It helps identify phosphorylated threonine motifs on kinase substrates, increasing binding affinity, while also providing a competitive site that DRA0333 may overrule under certain cellular contexts (Fig. 4). Studies on the catalytic residues D394 and D419, which are crucial for Mn²□ coordination, have confirmed that DRA0334’s phosphatase activity is depends on metal coordinating residues conserved for the PP2C family and dephosphorylation of RqkA and other STPKs (Fig. 4 & 5). Interestingly, the D394A mutant, despite being catalytically inactive, enhanced RqkA autophosphorylation (Fig. 4A). This suggests that even without functional PP2C domain, DRA0334’s KI-FHA module can act as canonical FHA domain module and stimulate the kinases phosphorylation (Fig. 4A & 5). This dual functionality possibly highlights the possible flexibility of FHA-like motifs, allowing them to shift between supportive and inhibitory roles based on their structural context and catalytic coupling. The competitive inhibition of phosphatase activity of DRA0334 by DRA0333 is demonstrated through both in vivo (Fig. 3) and in vitro (Fig. 4 & 5) experiments. This finding provides insight into FHA domain interaction with phosphatase suppression via the KI-FHA domain during specific regulation of RqkA and other STPKs. Notably, the KI-FHA domain of the DRA0334 phosphatase specifically targets the RqkA kinase, leading to enhanced kinase activity compared with other tested STPKs (DRA0332 and DR1243) (Fig. 4A, B). In contrast, when DRA0334 functions in combination with the FHA-domain protein DRA0333 tested, it appears to engage in broader kinase–phosphatase dynamic interactions. This highlights the functional significance of the operonic organization of the STPK (DRA0332), FHA-domain protein (DRA0333), and PP2C phosphatase (DRA0334) (Fig. 4 and 5).

Previously, the dynamic interaction of FHA domain proteins (GarA) and PknB STPK of *M. tuberculosis* shown to be modulated by phosphorylation of GarA by PknB STPK to induce intramolecular protein closure and dissociation of GarA from PknB kinase [73]. A similar type of regulation appears in this study’s data, where DRA0332 STPK is able to phosphorylate the FHA domain protein (DRA0333) (Fig. 4B & 5B). However, the effects of DRA0333 phosphorylation have not been tested here and will be addressed independently.

Additionally, an experiments with DRA0334 variants demonstrate that substrate docking and catalysis are related but separate processes (Fig. 5). A PP2C-only construct maintained highest catalytic phosphatase activity but lacked specificity. In contrast, the KI-FHA domain provided substrate selectivity and sensitivity to DRA0333’s competitive inhibition. A chimeric construct, which replaced DRA0334’s KI-FHA with DRA0333’s typical FHA, partially restored activity and improved substrate dephosphorylation under certain conditions (Fig. 5). This suggests that FHA domains can modulate the functions of both kinase and phosphatase when positioned correctly with regard to catalytic domains. This domain flexibility shows how FHA domain mediated phosphothreonine recognition and catalysis can be mixed to create dynamic signaling systems of context specific dynamic phosphorylation and dephosphorylation.

Moreover, DRA0334 has demonstrated a wide range of substrate specificity for the dephosphorylation of multiple STPKs of *D. radiodurans* (Fig. 3), suggesting it may acts as a central phosphatase hub that coordinates different phosphorylation networks in *D. radiodurans*. This role could have regulatory implications for a bacterium that needs to manage DNA repair, replication, and antioxidant responses while recovering from significant genomic damage. STRING-based interaction analyses back this idea by showing strong predicted links between DRA0334, RqkA, and other kinases related to DNA repair (Fig. S3 B). Considering the RqkA roles in DNA repair and cell division regulation [26, 34, 36, 41, 42, 65], findings of this study may indicate that the cycles of phosphorylation and dephosphorylation are closely tied to the DNA damage response. In terms of mechanics, though this study does not explore it, but it is tempting to suggest that the activation of kinase like RqkA likely drives repair and cell cycle arrest. Later, dephosphorylation by DRA0334 or a related STPP supports recovery once genome integrity is restored. This assumption is supported by previously published data showing that DRA0333 and its operonic partner are strongly overexpressed in an OxyR mutant, a newly identified oxidative stress sensor and transcriptional regulator activated under oxidative stress conditions [48]. However, comprehensive in vivo studies are still required to definitively establish the roles of the DRA0334 phosphatase and its operonic partner in the DNA damage response, which would be best addressed in a dedicated follow-up study.

Together, the present study uncovers a novel push–pull regulatory mechanism of phosphorylation in bacterial signal transduction. In this system, proteins containing operon-encoded manage serine/threonine phosphorylation through competition between kinase-interacting (KI-FHA) and standard phosphothreonine-binding FHA modules (Fig. 6). The model suggests that alone FHA-domain protein (DRA0333) or its divergent form KI-FHA domain typically support specific STPK functions by acting as phospho-dependent docking and regulatory modules (supported by data presented in Fig. 4). These module facilitate the kinase oligomerization, stabilize active conformations, and relieve autoinhibitory constraints to promote auto- and transphosphorylation. To counter or fine-tune the FHA domain’s effect on kinase signaling, PP2C-type phosphatases (DRA0334) utilize a divergent KI-FHA domain, allowing the dephosphorylation of kinases (RqkA / DRA0332) or phospho-substrates (RecA) to be adjusted according to signaling requirements (supported by data presented in Fig. 5). Thus, the proposed model reveals a previously unrecognized regulatory strategy in which both kinases and phosphatases employ FHA domain–mediated phosphosite docking to fine-tune their catalytic activities. This dynamic push–pull interplay between STPKs and STPPs likely coordinate signal transduction in *D. radiodurans* during recovery from acute genotoxic stress (Fig. 6). The proposed model is fundamentally different from the canonical linear phosphorylation cascades typically seen in eukaryotic systems. Instead, it shows how bacteria may have evolved a more modular approach to fine-tune the phospho-signaling for precise response.

Although biochemical and functional assays support FHA/KI-FHA–dependent regulation of STPK activity, the structural basis of these interactions remains unresolved. The proposed FHA–KI-FHA competition and operon-level docking model are inferred from functional data and will require high-resolution structural and in vivo validation. Future operon-level knockouts and stress-induction studies in *D. radiodurans* are needed to establish physiological relevance. Nevertheless, this work reinforces PP2C phosphatases as active regulators of phospho-signaling rather than passive terminators and identifies the DRA0331–DRA0334 operon as a modular, self-contained phosphorylation control unit integrating kinase activation, phosphatase feedback, and substrate recognition.

## Acknowledgments

Authors acknowledge Ms. Tarushi Singhal (SRF, HBNI) for her assistance and valuable suggestions in creating the homology model. Sincere thanks also go to Ms. Ankana Saha (SRF, HBNI) for her help in generating circular dichroism (CD) data. The authors express their gratitude to Dr. Hema Rajaram, Head of the Molecular Biology Division, and Dr. P.A. Hasan, Associate Director of the Biosciences Group at BARC, for their support and encouragement throughout this work. The author further acknowledges HBNI and BARC for providing financial support during this research.

## Material & Methods

### Bacterial Strains, growth media, antibodies, chemicals, and plasmids

For cloning, *Escherichia coli* NovaBlue was cultured in Luria–Bertani (LB) medium containing 1% NaCl, 1% tryptone, and 0.5% yeast extract at 37°C. The *E. coli* BTH101 strain (*cyaA*-deficient) was employed for *in vivo* protein–protein interaction analysis using the bacterial two-hybrid (BACTH) system and maintained at 30°C. Recombinant protein expression was carried out in *E. coli* BL21 (DE3) cells. Plasmids pET28a+, pUT18, pKNT25, and pRAD, along with their derivatives, were propagated in *E. coli* NovaBlue in the presence of the appropriate antibiotics. Standard molecular biology protocols were followed as previously described by Sambrook and Russell (2012) lab manual.

Antibodies specific to phospho-threonine/tyrosine residues (Cat. No. 9381S; Cell Signaling Technology) and the T18 domain of *Bordetella pertussis* CyaA protein (Cat. No. SC-13582; Santa Cruz Biotechnology) were used for immunoblotting. The anti-His antibody was obtained from New England BioLabs (NEB). All molecular-biology-grade reagents, enzymes, and salts were procured from Sigma-Aldrich, Roche Biochemicals (Mannheim, Germany), NEB, and Merck India Pvt. Ltd. Radiolabeled nucleotides [³²P] were supplied by the Board of Radiation and Isotope Technology (BRIT), Department of Atomic Energy, India.

### Recombinant plasmid construction and protein purification

Table 1 has a list of plasmids and primers used in this study. The transnational fused DRA0334, DR2513, DR1243 with T18 or T25-tag at generated in the pUT18 and pKNT25 vectors plasmids. Plasmids were transformed into BTH101 *E. coli* cells in different combination and permutations to see in vivo protein-protein interaction as describe elsewhere [65]. The expression of fusion proteins in BTH101 cells was confirmed by Western blotting using antibodies specific to the T18 / T25 tags.

For protein purification, genes encoding the proteins listed in Table 1 were cloned into the pET28a expression vector to generate N-terminal His-tagged constructs, which were transformed into *E. coli* BL21 (DE3) cells for overexpression. One litre cultures were grown in LB medium containing kanamycin (50 µg/mL) at 37 °C to an OD□□□ of 0.6–0.8 and induced with 500 µM IPTG, followed by further incubation for 4–6 h at 30 °C (or overnight at 18 °C where required). Cells were harvested by centrifugation and resuspended in buffer A (20 mM Tris-HCl, pH 7.6; 300 mM NaCl; 10% glycerol) supplemented with 0.5 mg/mL lysozyme, 1 mM PMSF, and 0.02% Triton X-100, and incubated at 37 °C for 30 min, after which a protease inhibitor cocktail (NEB) was added. Cell lysis was completed by sonication on ice for 10 min using 5-second pulses with 10-second cooling intervals at 35% amplitude, and the lysate was clarified by centrifugation at 12,000 rpm for 30 min at 4 °C. The supernatant was applied to a Ni²□-charged chelating Sepharose column (GE Healthcare) pre-equilibrated with buffer A, washed with 20 column volumes of buffer A containing 20 mM imidazole, and the bound proteins were eluted with buffer A supplemented with 300 mM imidazole. Eluted fractions were analyzed by SDS–PAGE, pooled, and further purified by gel-filtration chromatography on a Superdex 200 column using buffer B (20 mM Tris-HCl, pH 7.6, 1M NaCl, 1mM DTT, 1mM EDTA, and 10% glycerol), and fractions exhibiting >95% purity were concentrated by ammonium sulphate precipitation, dialyzed against buffer C (10 mM Tris-HCl, pH 7.6, 50 mM NaCl, 1mM DTT, 1mM EDTA, 50% glycerol, 1 mM PMSF), aliquoted, and stored at −20 °C.

### Bacterial two-hybrid assay (Protein–protein interaction analysis)

*In vivo* protein–protein interactions were examined using the BACTH system as described by Sharma *et al.* [74]. BTH101 cells were co-transformed with different pUT18 and pKNT25 constructs of *drA0334, dr1243, dr2513,* and *rqkA* expressing target proteins fused with T18 or T25 tags at the C-terminus in different combinations. Empty vectors (pUT18 and pKNT25) served as negative controls. Transformants were plated on LB agar supplemented with 40 µg/mL 5-bromo-4-chloro-3-indolyl-β-d-galactopyranoside (X-Gal), 0.5 mM IPTG, and appropriate antibiotics, followed by incubation at 30°C for 12 h. Protein–protein interactions were visualized based on blue/white colony development.

### Cloning and site-directed mutagenesis

Genomic DNA from *D. radiodurans* was isolated using a commercial kit (BRIT, DAE, India). The *drA0334* gene was amplified using gene-specific primers (Table 1) and cloned into *pET28a* at *EcoRI* and *XhoI* restriction sites, yielding plasmid pET28a-*drA0334*. Site-directed mutagenesis of conserved residues Asp394 and Asp419 was performed by two-step overlapping PCR. All constructs were confirmed by sequencing and subsequently transformed into *E. coli* BL21 (DE3) cells for recombinant protein expression.

### Biochemical characterization

The phosphatase activity of DRA0334 was assessed using p-nitrophenyl phosphate (PNPP) as the substrate. Reaction conditions were initially optimized for pH (7.6 and 8.8) and divalent metal ion dependence using 50 nM DRA0334 and 500 μM PNPP in buffer D (20 mM Tris–HCl, pH 7.6, 100 mM NaCl, and 1 mM divalent metal ions). Phosphate release was quantified as nanomoles of p-nitrophenol (PNP) per microgram of protein, calculated using an extinction coefficient at pH 7.6 of 13286 M□¹·cm□¹. Enzyme kinetics were determined using Michaelis–Menten and Lineweaver–Burk analyses. For kinetic assays, reactions were performed in triplicate in 96-well plates using buffer D supplemented with 1 mM Mn²□, containing 50 nM DRA0334 or its mutant variants and increasing concentrations of PNPP (0.5–50 mM). Absorbance was monitored continuously without NaOH termination to prevent precipitation. Km and Vmax values were obtained by fitting the data to the Michaelis–Menten equation, and Lineweaver–Burk plots were generated using GraphPad Prism.

### Ex vivo dephosphorylation assay

The ex vivo dephosphorylation assayed in *E. coli* BL21 (DE3) or NovaBlue strains. Plasmids pET28a-*drA0334* and pET28a-*dr*2513 were co-transformed with pRAD-RqkA or pRAD-1243 into *E. coli* BL21 (DE3) for inducible expression of DRA0334 and DR2513 STPPs and constitutive expression of RqkA or DR1243 STPKs. While for constitutive expression DRA0334 and DR2513 STPPs and RqkA or DR1243 STPKs, plasmids *drA0334* pKNT25 and *dr2513* pKNT25 were co-transformed with pRAD-RqkA or pRAD-1243 into for NovaBlue strains. Protein expression was induced in BL21 with IPTG, while NovaBlue cells exhibited constitutive expression. Equivalent cell numbers were lysed and analyzed by SDS–PAGE followed by western blotting using anti-phospho-threonine antibody.

### In vitro dephosphorylation assay

To generate phosphorylated STPKs (RqkA, DRA0332, and DR1243), purified kinases (5–10 µM) were incubated in PNK buffer (50 mM Tris–HCl, 10 mM MgCl□, 5 mM DTT, and 0.1 mM ATP) supplemented with [³²P]-γ-ATP for 30 min at 37 °C. Phosphorylated RecA and MBP were obtained by incubating each substrate (10 µM) individually with RqkA (1 µM) in the same buffer, maintaining a substrate-to-kinase molar ratio of 10:1, for 30 min at 37 °C. Excess [³²P]-γ-ATP was removed by passage through a G-50 Sephadex column, and the resulting phosphorylated STPKs or phosphoproteins were subsequently used as substrates for dephosphorylation assays. Dephosphorylation reactions were performed using DRA0334 phosphatase at a phosphatase-to-substrate molar ratio of 1:3 in buffer D (20 mM Tris–HCl, pH 7.6, 100 mM NaCl) supplemented with 1 mM Mn²□ and incubated for 1 h. Reactions were terminated, resolved by SDS–PAGE, and the gels were dried and exposed to X-ray film for autoradiography.

### Biophysical characterization

Protein folding and structural integrity were assessed by circular dichroism (CD) spectroscopy. CD measurements were performed using a J-815 spectropolarimeter (JASCO, Japan) at room temperature. Purified protein samples were prepared in 10 mM Tris–HCl (pH 7.6) containing 10 mM NaCl and equilibrated prior to data acquisition. Far-UV CD spectra were recorded over the wavelength range of 190–260 nm using a quartz cuvette with a 1-mm path length. Spectra were collected with a bandwidth of 1 nm, a scanning speed of 50 nm/min, and a response time of 1 s. Each spectrum represents the average of three consecutive scans, and the corresponding buffer spectrum was recorded under identical conditions and subtracted from the sample spectra. The resulting CD data were expressed as mean residue ellipticity and used to evaluate secondary structure content and overall protein folding.

### Homology modelling

Homology modeling of *D. radiodurans* PP2C-type phosphatases used AlphaFold and lab-determined structural templates to study domain conservation and the arrangement of catalytic sites. The AlphaFold models for DR2513 (AF-Q9RRH8-F1-model_v6.pdb) and DRA0334 (AF-Q9RYH8-F1-model_v6.pdb) were utilized alongside the crystallographic structure of *M. tuberculosi*s PstP (PDB ID: 1TXO) as a reference. To define the conserved phosphatase core region, sequence alignment was done using amino acid residues 10 to 244 of DR2513, 376 to 616 of DRA0334, and 9 to 238 of PstP.

## Data Availability statement

The authors confirm that the data supporting the findings of this study are available within the article or its supplementary materials. However, specific data that support the findings of this study are available from the corresponding author, upon request.

## References

1. Pereira, S.F., L. Goss, and J. Dworkin, Eukaryote-like serine/threonine kinases and phosphatases in bacteria. Microbiol Mol Biol Rev, 2011. 75(1): p. 192–212.

2. Prisic, S. and R.N. Husson, Mycobacterium tuberculosis Serine/Threonine Protein Kinases. Microbiol Spectr, 2014. 2(5).

3. Zeng, J., et al., Protein kinases PknA and PknB independently and coordinately regulate essential Mycobacterium tuberculosis physiologies and antimicrobial susceptibility. PLoS Pathog, 2020. 16(4): p. e1008452.

4. Hardt, P., et al., The cell wall precursor lipid II acts as a molecular signal for the Ser/Thr kinase PknB of Staphylococcus aureus. Int J Med Microbiol, 2017. 307(1): p. 1–10.

5. Garcia, P.S., et al., Cell division of Streptococcus pneumoniae: think positive! Curr Opin Microbiol, 2016. 34: p. 18–23.

6. Fiuza, M., et al., From the characterization of the four serine/threonine protein kinases (PknA/B/G/L) of Corynebacterium glutamicum toward the role of PknA and PknB in cell division. J Biol Chem, 2008. 283(26): p. 18099–112.

7. Gaidenko, T.A., T.J. Kim, and C.W. Price, The PrpC serine-threonine phosphatase and PrkC kinase have opposing physiological roles in stationary-phase Bacillus subtilis cells. J Bacteriol, 2002. 184(22): p. 6109–14.

8. Sharma, A.K., et al., Serine/Threonine Protein Phosphatase PstP of Mycobacterium tuberculosis Is Necessary for Accurate Cell Division and Survival of Pathogen. J Biol Chem, 2016. 291(46): p. 24215–24230.

9. Jarick, M., et al., The serine/threonine kinase Stk and the phosphatase Stp regulate cell wall synthesis in Staphylococcus aureus. Sci Rep, 2018. 8(1): p. 13693.

10. Osaki, M., et al., The StkP/PhpP signaling couple in Streptococcus pneumoniae: cellular organization and physiological characterization. J Bacteriol, 2009. 191(15): p. 4943–50.

11. Wagner, M.J., et al., Molecular mechanisms of SH2- and PTB-domain-containing proteins in receptor tyrosine kinase signaling. Cold Spring Harb Perspect Biol, 2013. 5(12): p. a008987.

12. Bradley, D. and P. Beltrao, Evolution of protein kinase substrate recognition at the active site. PLoS Biol, 2019. 17(6): p. e3000341.

13. Yaffe, M.B. and A.E. Elia, Phosphoserine/threonine-binding domains. Curr Opin Cell Biol, 2001. 13(2): p. 131–8.

14. Hofmann, K. and P. Bucher, The FHA domain: a putative nuclear signalling domain found in protein kinases and transcription factors. Trends Biochem Sci, 1995. 20(9): p. 347–9.

15. Almawi, A.W., L.A. Matthews, and A. Guarne, FHA domains: Phosphopeptide binding and beyond. Prog Biophys Mol Biol, 2017. 127: p. 105–110.

16. Liang, X. and S.R. Van Doren, Mechanistic insights into phosphoprotein-binding FHA domains. Acc Chem Res, 2008. 41(8): p. 991–9.

17. Sun, Z., et al., Rad53 FHA domain associated with phosphorylated Rad9 in the DNA damage checkpoint. Science, 1998. 281(5374): p. 272-4.

18. Durocher, D. and S.P. Jackson, The FHA domain. FEBS Lett, 2002. 513(1): p. 58–66.

19. Ahn, J.Y., et al., Phosphorylation of threonine 68 promotes oligomerization and autophosphorylation of the Chk2 protein kinase via the forkhead-associated domain. J Biol Chem, 2002. 277(22): p. 19389–95.

20. Kim, K., et al., Phosphopeptide interactions of the Nbs1 N-terminal FHA-BRCT1/2 domains. Sci Rep, 2021. 11(1): p. 9046.

21. Wang, Q., The role of forkhead-associated (FHA)-domain proteins in plant biology. Plant Mol Biol, 2023. 111(6): p. 455–472.

22. Wang, G., et al., Type VI secretion system-associated FHA domain protein TagH regulates the hemolytic activity and virulence of Vibrio cholerae. Gut Microbes, 2022. 14(1): p. 2055440.

23. Mougous, J.D., et al., Threonine phosphorylation post-translationally regulates protein secretion in Pseudomonas aeruginosa. Nat Cell Biol, 2007. 9(7): p. 797–803.

24. Daly, M.J., et al., Protein oxidation implicated as the primary determinant of bacterial radioresistance. PLoS Biol, 2007. 5(4): p. e92.

25. Misra, H.S., Y.S. Rajpurohit, and N.P. Khairnar, Pyrroloquinoline-quinone and its versatile roles in biological processes. J Biosci, 2012. 37(2): p. 313–25.

26. Rajpurohit, Y.S., S.S. Desai, and H.S. Misra, Pyrroloquinoline quinone and a quinoprotein kinase support gamma-radiation resistance in Deinococcus radiodurans and regulate gene expression. J Basic Microbiol, 2013. 53(6): p. 518–31.

27. Rajpurohit, Y.S., R. Gopalakrishnan, and H.S. Misra, Involvement of a protein kinase activity inducer in DNA double strand break repair and radioresistance of Deinococcus radiodurans. J Bacteriol, 2008. 190(11): p. 3948–54.

28. Zahradka, K., et al., Reassembly of shattered chromosomes in Deinococcus radiodurans. Nature, 2006. 443(7111): p. 569–73.

29. Qi, H.Z., et al., Antioxidative system of Deinococcus radiodurans. Res Microbiol, 2020. 171(2): p. 45–54.

30. Xu, G., et al., DdrB stimulates single-stranded DNA annealing and facilitates RecA-independent DNA repair in Deinococcus radiodurans. DNA Repair (Amst), 2010. 9(7): p. 805–12.

31. Slade, D., et al., Recombination and replication in DNA repair of heavily irradiated Deinococcus radiodurans. Cell, 2009. 136(6): p. 1044–55.

32. Sharma, D.K., et al., Natural transformation-specific DprA coordinate DNA double-strand break repair pathways in heavily irradiated D. radiodurans. Appl Environ Microbiol, 2024. 90(2): p. e0194823.

33. Bouthier de la Tour, C., et al., The deinococcal DdrB protein is involved in an early step of DNA double strand break repair and in plasmid transformation through its single-strand annealing activity. DNA Repair (Amst), 2011. 10(12): p. 1223–31.

34. Misra, H.S. and Y.S. Rajpurohit, DNA damage response and cell cycle regulation in bacteria: a twist around the paradigm. Front Microbiol, 2024. 15: p. 1389074.

35. Chaudhary, R., et al., FtsZ phosphorylation brings about growth arrest upon DNA damage in Deinococcus radiodurans. FASEB Bioadv, 2023. 5(1): p. 27–42.

36. Rajpurohit, Y.S., D.K. Sharma, and H.S. Misra, Involvement of serine / threonine protein kinases in DNA damage response and cell division in bacteria. Res Microbiol, 2022. 173(1-2): p. 103883.

37. Sharma, D.K., et al., WD40 domain of RqkA regulates its kinase activity and role in extraordinary radioresistance of D. radiodurans. J Biomol Struct Dyn, 2022. 40(3): p. 1246–1259.

38. Sharma, D.K., et al., Phosphorylation of deinococcal RecA affects its structural and functional dynamics implicated for its roles in radioresistance of Deinococcus radiodurans. J Biomol Struct Dyn, 2020. 38(1): p. 114–123.

39. Maurya, G.K., et al., Phosphorylation of FtsZ and FtsA by a DNA Damage-Responsive Ser/Thr Protein Kinase Affects Their Functional Interactions in Deinococcus radiodurans. mSphere, 2018. 3(4).

40. Rajpurohit, Y.S., et al., Phosphorylation of Deinococcus radiodurans RecA Regulates Its Activity and May Contribute to Radioresistance. J Biol Chem, 2016. 291(32): p. 16672–85.

41. Rajpurohit, Y.S. and H.S. Misra, Structure-function study of deinococcal serine/threonine protein kinase implicates its kinase activity and DNA repair protein phosphorylation roles in radioresistance of Deinococcus radiodurans. Int J Biochem Cell Biol, 2013. 45(11): p. 2541–52.

42. Rajpurohit, Y.S. and H.S. Misra, Characterization of a DNA damage-inducible membrane protein kinase from Deinococcus radiodurans and its role in bacterial radioresistance and DNA strand break repair. Mol Microbiol, 2010. 77(6): p. 1470–82.

43. Sancar, A., et al., Molecular mechanisms of mammalian DNA repair and the DNA damage checkpoints. Annu Rev Biochem, 2004. 73: p. 39–85.

44. Makarova, K.S., et al., Genome of the extremely radiation-resistant bacterium Deinococcus radiodurans viewed from the perspective of comparative genomics. Microbiol Mol Biol Rev, 2001. 65(1): p. 44–79.

45. Pallen, M., R. Chaudhuri, and A. Khan, Bacterial FHA domains: neglected players in the phospho-threonine signalling game? Trends Microbiol, 2002. 10(12): p. 556–63.

46. Jiang, Z., et al., The FHA domain protein ArnA functions as a global DNA damage response repressor in the hyperthermophilic archaeon Saccharolobus islandicus. mBio, 2023. 14(4): p. e0094223.

47. Whittaker, C.A. and R.O. Hynes, Distribution and evolution of von Willebrand/integrin A domains: widely dispersed domains with roles in cell adhesion and elsewhere. Mol Biol Cell, 2002. 13(10): p. 3369–87.

48. Chen, H., et al., A novel OxyR sensor and regulator of hydrogen peroxide stress with one cysteine residue in Deinococcus radiodurans. PLoS One, 2008. 3(2): p. e1602.

49. Missiakas, D. and S. Raina, Signal transduction pathways in response to protein misfolding in the extracytoplasmic compartments of E. coli: role of two new phosphoprotein phosphatases PrpA and PrpB. EMBO J, 1997. 16(7): p. 1670–85.

50. Halbedel, S., et al., Regulatory protein phosphorylation in Mycoplasma pneumoniae. A PP2C-type phosphatase serves to dephosphorylate HPr(Ser-P). J Biol Chem, 2006. 281(36): p. 26253–9.

51. Schlicker, C., et al., Structural analysis of the PP2C phosphatase tPphA from Thermosynechococcus elongatus: a flexible flap subdomain controls access to the catalytic site. J Mol Biol, 2008. 376(2): p. 570–81.

52. Treuner-Lange, A., M.J. Ward, and D.R. Zusman, Pph1 from Myxococcus xanthus is a protein phosphatase involved in vegetative growth and development. Mol Microbiol, 2001. 40(1): p. 126–40.

53. Rajagopal, L., et al., Regulation of purine biosynthesis by a eukaryotic-type kinase in Streptococcus agalactiae. Mol Microbiol, 2005. 56(5): p. 1329–46.

54. Ulrych, A., et al., Characterization of pneumococcal Ser/Thr protein phosphatase phpP mutant and identification of a novel PhpP substrate, putative RNA binding protein Jag. BMC Microbiol, 2016. 16(1): p. 247.

55. Das, A.K., et al., Crystal structure of the protein serine/threonine phosphatase 2C at 2.0 A resolution. EMBO J, 1996. 15(24): p. 6798–809.

56. Bork, P., et al., The protein phosphatase 2C (PP2C) superfamily: detection of bacterial homologues. Protein Sci, 1996. 5(7): p. 1421–5.

57. Kamada, R., et al., Metal-dependent Ser/Thr protein phosphatase PPM family: Evolution, structures, diseases and inhibitors. Pharmacol Ther, 2020. 215: p. 107622.

58. Greenstein, A.E., et al., Structure/function studies of Ser/Thr and Tyr protein phosphorylation in Mycobacterium tuberculosis. J Mol Microbiol Biotechnol, 2005. 9(3-4): p. 167–81.

59. Rajagopalan, K., J. Dworkin, and E. Nagle, Identification and Biochemical Characterization of a Novel Protein Phosphatase 2C-Like Ser/Thr Phosphatase in Escherichia coli. J Bacteriol, 2018. 200(18).

60. Chopra, P., et al., Phosphoprotein phosphatase of Mycobacterium tuberculosis dephosphorylates serine-threonine kinases PknA and PknB. Biochem Biophys Res Commun, 2003. 311(1): p. 112–20.

61. Obuchowski, M., et al., Characterization of PrpC from Bacillus subtilis, a member of the PPM phosphatase family. J Bacteriol, 2000. 182(19): p. 5634–8.

62. Boitel, B., et al., PknB kinase activity is regulated by phosphorylation in two Thr residues and dephosphorylation by PstP, the cognate phospho-Ser/Thr phosphatase, in Mycobacterium tuberculosis. Mol Microbiol, 2003. 49(6): p. 1493–508.

63. Arora, G., et al., Understanding the role of PknJ in Mycobacterium tuberculosis: biochemical characterization and identification of novel substrate pyruvate kinase A. PLoS One, 2010. 5(5): p. e10772.

64. Novakova, L., et al., Characterization of a eukaryotic type serine/threonine protein kinase and protein phosphatase of Streptococcus pneumoniae and identification of kinase substrates. FEBS J, 2005. 272(5): p. 1243–54.

65. Rajpurohit, Y.S., D.K. Sharma, and H.S. Misra, PprA Protein Inhibits DNA Strand Exchange and ATP Hydrolysis of Deinococcus RecA and Regulates the Recombination in Gamma-Irradiated Cells. Front Cell Dev Biol, 2021. 9: p. 636178.

66. Li, J., G.P. Smith, and J.C. Walker, Kinase interaction domain of kinase-associated protein phosphatase, a phosphoprotein-binding domain. Proc Natl Acad Sci U S A, 1999. 96(14): p. 7821–6.

67. Niwa, S., T. Watanabe, and K. Chiba, The FHA domain is essential for autoinhibition of KIF1A/UNC-104 proteins. J Cell Sci, 2024. 137(19).

68. Reinhardt, H.C. and M.B. Yaffe, Phospho-Ser/Thr-binding domains: navigating the cell cycle and DNA damage response. Nat Rev Mol Cell Biol, 2013. 14(9): p. 563–80.

69. Spivey, V.L., et al., Forkhead-associated (FHA) domain containing ABC transporter Rv1747 is positively regulated by Ser/Thr phosphorylation in Mycobacterium tuberculosis. J Biol Chem, 2011. 286(29): p. 26198–209.

70. Molle, V., et al., Two FHA domains on an ABC transporter, Rv1747, mediate its phosphorylation by PknF, a Ser/Thr protein kinase from Mycobacterium tuberculosis. FEMS Microbiol Lett, 2004. 234(2): p. 215–23.

71. Trotochaud, A.E., et al., The CLAVATA1 receptor-like kinase requires CLAVATA3 for its assembly into a signaling complex that includes KAPP and a Rho-related protein. Plant Cell, 1999. 11(3): p. 393–406.

72. Braun, D.M., J.M. Stone, and J.C. Walker, Interaction of the maize and Arabidopsis kinase interaction domains with a subset of receptor-like protein kinases: implications for transmembrane signaling in plants. Plant J, 1997. 12(1): p. 83–95.

73. England, P., et al., The FHA-containing protein GarA acts as a phosphorylation-dependent molecular switch in mycobacterial signaling. FEBS Lett, 2009. 583(2): p. 301–7.

74. Sharma, D.K., et al., Characterization of DNA Processing Protein A (DprA) of the Radiation-Resistant Bacterium Deinococcus radiodurans. Microbiol Spectr, 2022. 10(6): p. e0347022.

