## Supplementary Data file for "Operon-driven antagonism and novel modulation of DNA damage sensor STPK specific PP2C phosphatase in *Deinococcus radiodurans*"

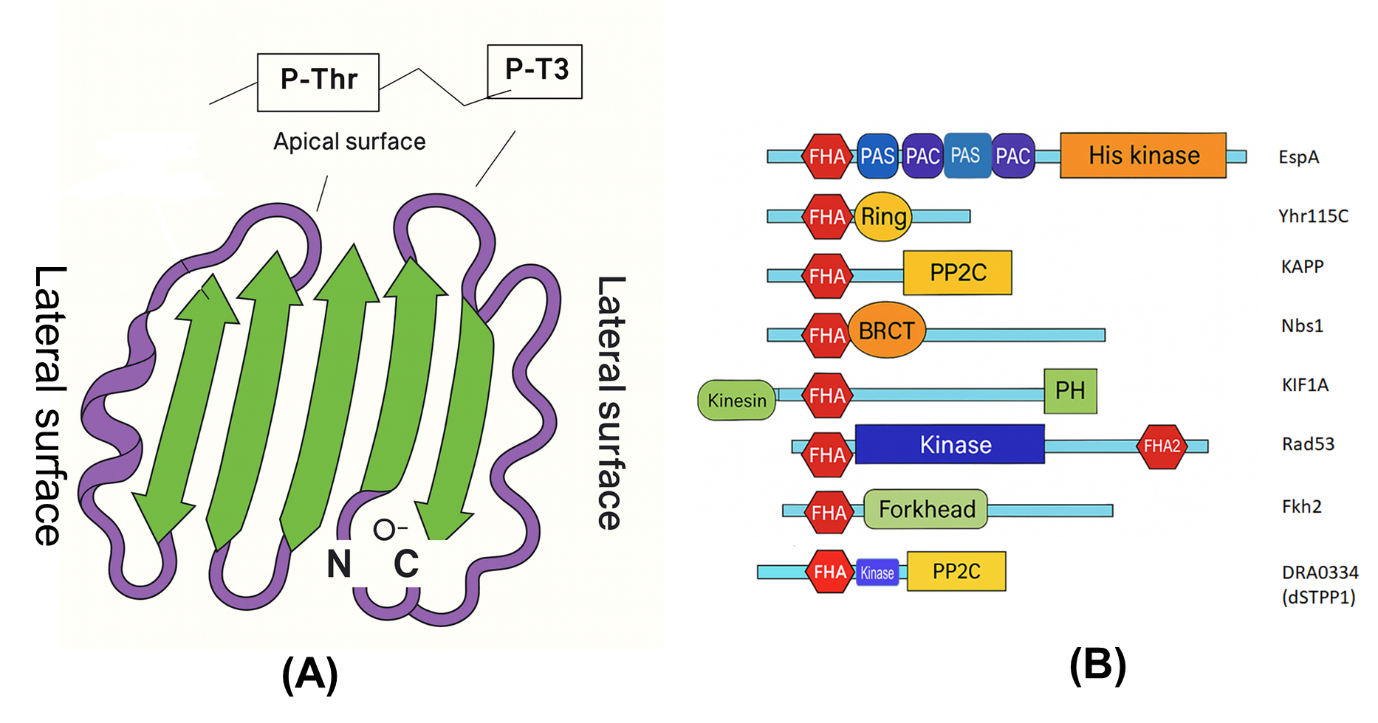
**Figure S1. Structural and domain diversity of FHA domains proteins.** (A) Structural model of FHA domain in interaction with phosphor-threonine residue. (B) Modular organization of FHA domain with variety of other functional domains. Phosphatase domain (PP2C), BRCT, and kinase domain abundantly associated with FHA domain.

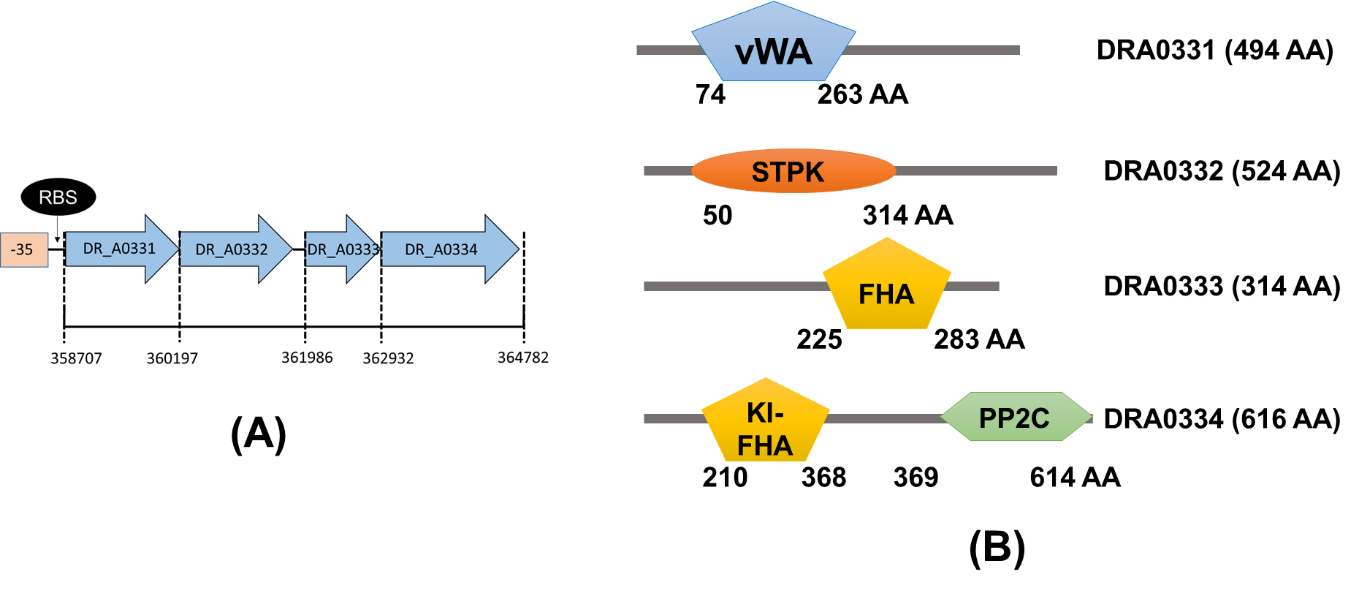

**Figure S2. Operon organisation and domains of key operon proteins.** (A) Operon organisation of DRA0331 to DRA0334 proteins. (B) Domain architecture of operon partner proteins showing the protein length in number of amino acids a start and end boundary of different domains in individual protein.

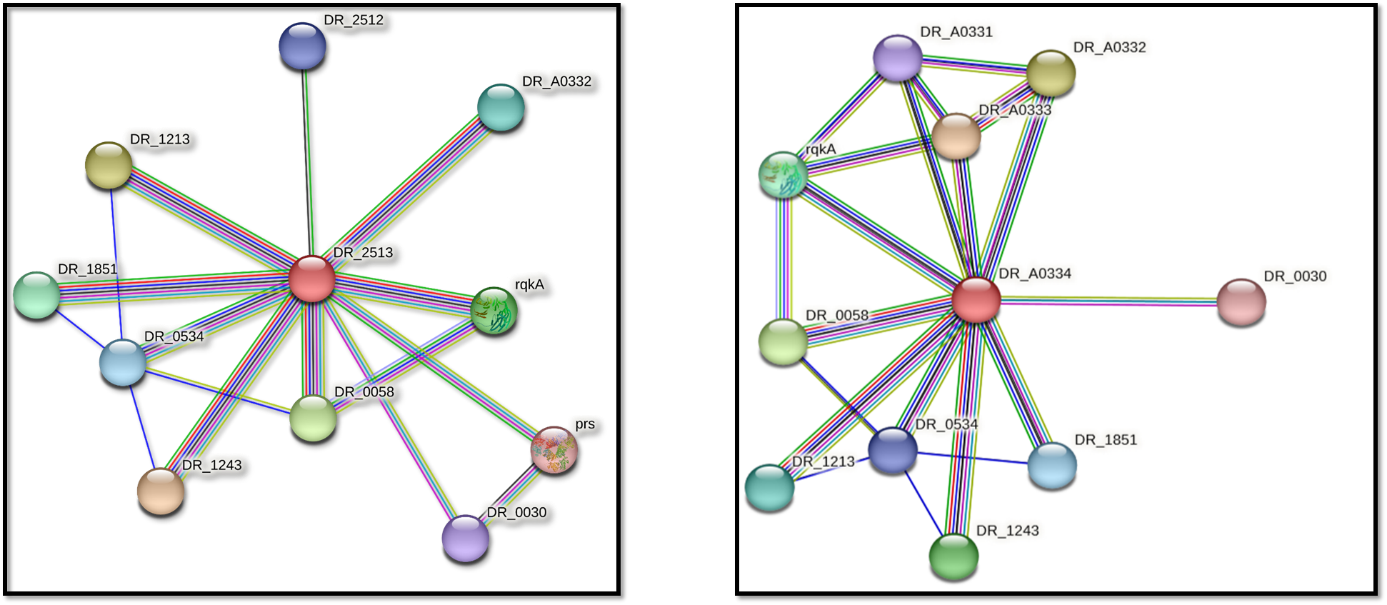

**Figure S3. Predicted interaction network of DRA0334 and DR2513 PP2C phosphatases.** STRING analysis illustrating strong associations of DRA0334 and DR2513 with several STPKs and DNA repair proteins, suggesting coordinated regulation of phosphorylation–dephosphorylation pathways.

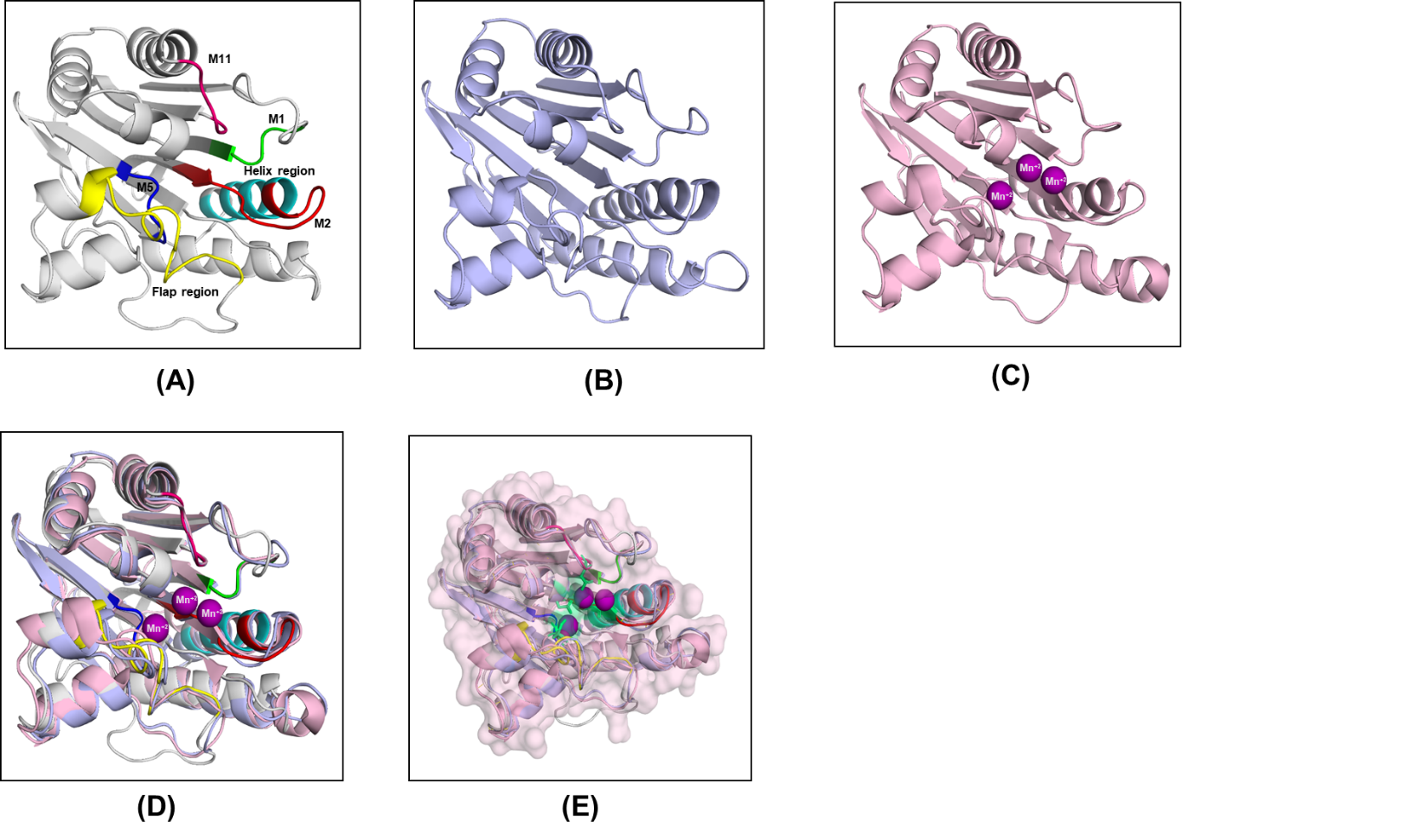

**Figure S4. Structural modeling of *D. radiodurans* PP2C phosphatases.** Structural models of *Deinococcus radiodurans* phosphatases DR2513 and DRA0334 and *Mycobacterium tuberculosis* PstP were analyzed to compare conserved PP2C catalytic features. (A) the ribbon model of DRA0334 is shown in grey with key motifs highlighted: M1 (green), M2 (red), M5 (blue), M11 (hot pink), flap region (yellow), and helix region (cyan). (B) depicts DR2513 as a blue chain, while (C) shows PstP as a pink chain with Mn²⁺ ions represented in magenta. (D) presents a superimposed ribbon model of DR2513 and DRA0334 phosphatase domains overlaid on PstP, illustrating conserved structural folds. (E) displays a superimposed view showing only the active-site aspartate residues of PstP as green helix and Mn²⁺ ions represented in magenta, aligned with ribbon representations of DR2513 and DRA0334 phosphatase domains, highlighting active-site conservation and catalytic geometry.

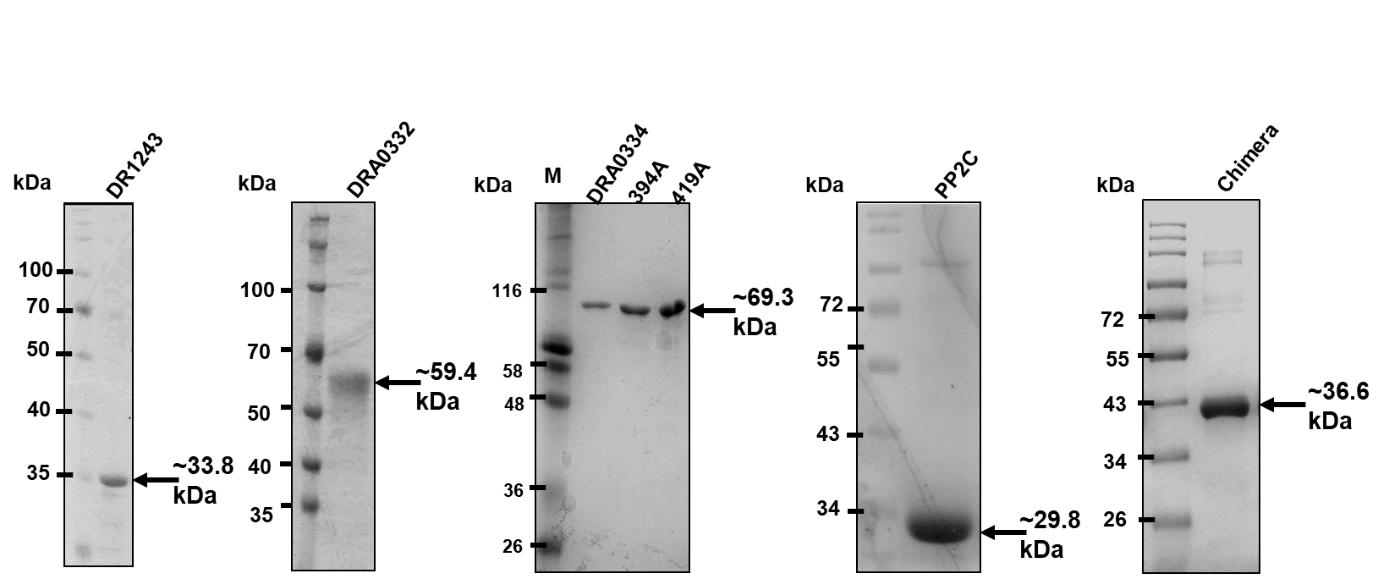

**Figure S5. Purification of DRA0334 and catalytic mutants.** SDS–PAGE analysis of purified wild-type DRA0334 and its D394A, D419A, PP2C-only, and chimera variants used for biochemical assays. Additionally, purified DR1243 and DRA0332 STPKs shown.

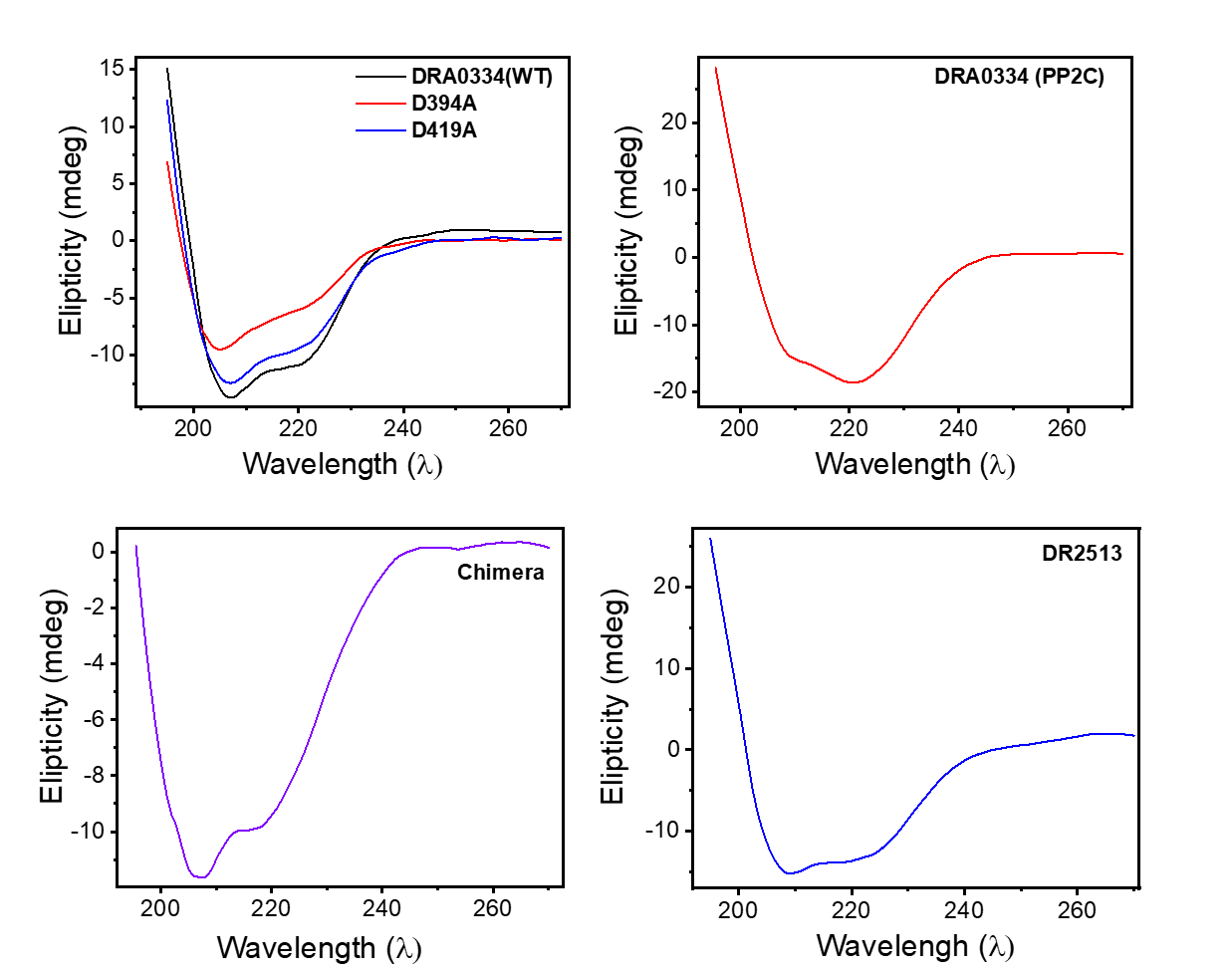

**Figure S6. Secondary-structure integrity of DRA0334, its mutants, and DR2513 PP2C phosphatase.** Circular dichroism spectra showing that alanine substitutions at catalytic residues or chimera do not perturb the overall protein fold. While PP2C-only domain showed shift in alpha-helical conformation compare to DRA0334 wild-type.

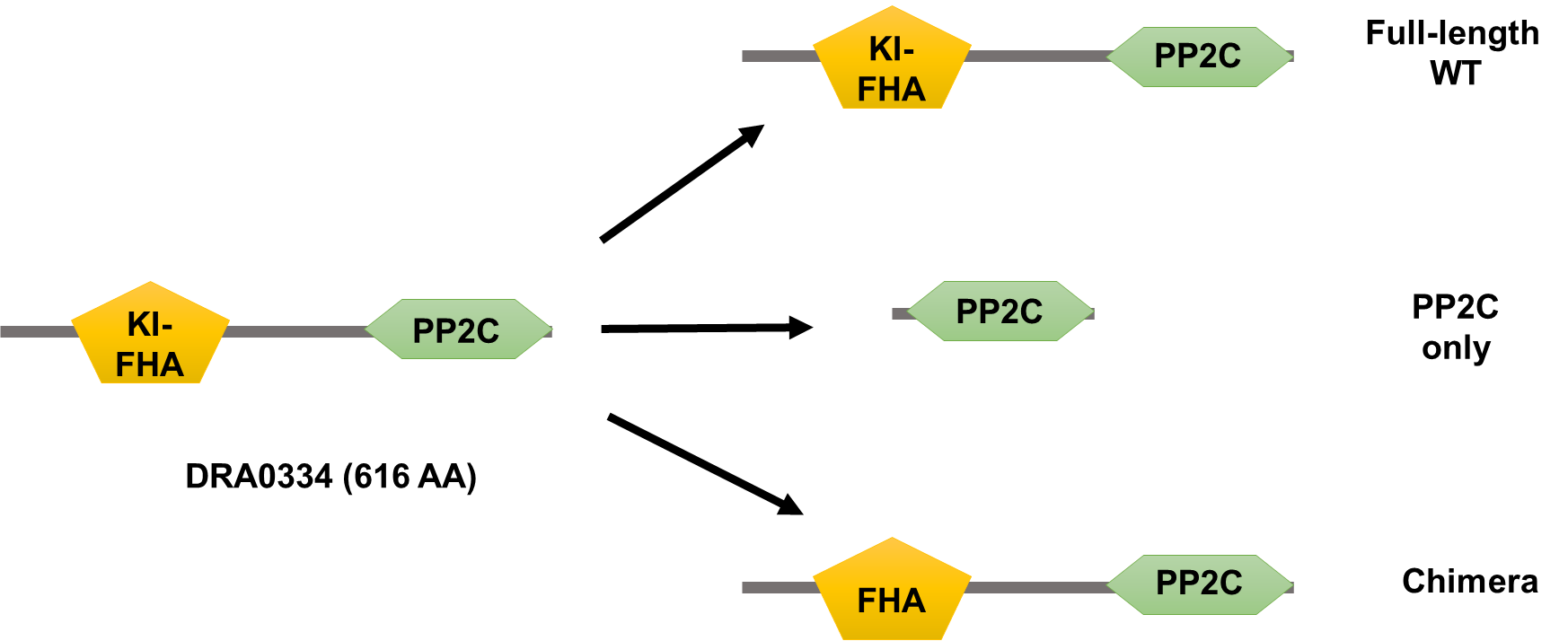

**Figure S7. Schematic of DRA0334 constructs used in this study.** Diagram showing domain organization of full-length DRA0334, PP2C-only, and FHA-chimeric variants used for functional characterization of phosphatase activity.

**Table S1: Bacterial strains, plasmids, and primers Used in this study**

**A. Bacterial Strains**

| **Bacterial strains** | **Organism** | **Genotype** | **Source** |
| --- | --- | --- | --- |
| *D. radiodurans* R1 | *D. radiodurans* | Wild type strain ATCC13939 | Lab stock |
| *E. coli* Novablue | *E. coli* | *end*A1 *hsd*R17*(r _K12_ ^−^ m _K12_ ^+^) sup*E44 *thi-1 rec*A1 *gyr*A96 *rel*A1 *lac*F’'*[pro*A^+^B*^+^ lac*I^q^ *Z∆*M15*::*T*n*10] (Tet^R^) | Lab stock |
| *E. coli* BTH 101 | *E. coli* | F^-^, *cya-*99*, ara*D139*, gal*E15*, gal*K16*, rps*L1 (Str ^r^)*, hsd*R2*, mcr*A1*, mcr*B1 | Lab stock |
| *E. coli Top10* | *E. coli* | F- mcrA Δ(mrr-hsdRMS-mcrBC) Φ80lacZΔM15 Δ lacX74 recA1 araD139 Δ( araleu)7697 galU galK rpsL (StrR) endA1 nupG | Lab stock |
| *E. coli* BL21(DE3) | *E. coli* | *fhu*A2 *(lon)omp*T *gal*(λDE3)*(dcm) ∆hs*dS | Lab stock |

**Plasmids:**

| **Names** | **Characteristics and Source** | lab stock |
| --- | --- | --- |
| pVHS559 | A shuttle vector between *D. radiodurans* and *E. coli* (Spec^R^) | lab stock |
| pRADgro | pRAD1 carrying 261bp *Bgl*II-*Xba*I fragment of promoter (Pgro) from *D. radiodurans* | lab stock |
| pUT18 | pUC19 derivative, MCS at N-terminal of T18 fragments of adenylate cyclase, ~3 kb, AmpR | lab stock |
| pKNT25 | pSU40 derivative, MCS at N-terminal of T25 fragment of adenylate cyclase, ~3.4 kb, KanR | lab stock |
| *rqkA* pKNT25/pUT18 | pKNT25 or pUT18 vectors carrying *dr2518* gene | Lab Stock |
| pET28a-*rqkA* | pET28a vector carrying *dr2518* gene | Lab Stock |
| pRAD-*rqkA* | pRAD vector carrying *dr2518* gene | Lab Stock |
| pET28a-*drA0334* | pET28a vector carrying *drA0334* gene at EcoR1 and XhoI | This work |
| pET28a-*dr2513* | pET28a vector carrying *dr2513* gene at EcoR1 and XhoI | This work |
| pET28a-*dr1243* | pET28a vector carrying *dr1243* gene at EcoR1 and XhoI | This work |
| pET28a-*drA0332* | pET28a vector carrying *drA0332* gene at BamHI and HindIII | This work |
| pET28a-*PP2C* | pET28a vector carrying PP2C domain of *drA0334* gene at EcoR1 and XhoI | This work |
| pET28a-*D394A* | pET28a vector carrying *D394A mutant of drA0334* gene at EcoR1 and XhoI | This work |
| pET28a-*D419A* | pET28a vector carrying *D419A mutant of drA0334* gene at EcoR1 and XhoI | This work |
| pET28a-*drA0333* | pET28a vector carrying *drA0333* gene at EcoR1 and XhoI | This work |
| pET28a-*Chimera* | pET28a vector carrying Chimera (FHA domain of DRA0333 and PP2C domain of DRA0334) at EcoR1 and XhoI | This work |
| pRAD-*dr1243* | pRAD vector carrying *dr1243* gene at ApaI and XbaI | This work |
| *drA0334* pKNT25/pUT18 | pKNT25 or pUT18 vectors carrying *drA0334* gene at XbaI and KpnI | This work |
| *dr2513* pKNT25/pUT18 | pKNT25 or pUT18 vectors carrying *dr2513* gene at XbaI and KpnI | This work |
| *dr1243* pKNT25/pUT18 | pKNT25 or pUT18 vectors carrying *dr1243* gene at XbaI and KpnI | This work |

**Primers used:**

| **SI. No.** | **Name of primer and** | **Sequence (5’ to 3’)** | **Purpose** |
| --- | --- | --- | --- |
| 1 | pET28a-*drA0334* | GGAATTCCATATGGTGAACGAGAACCACCCGCG (F)  CCGCTCGAGTCACGCCGTTCTCTCGATCACCAG (R) | Cloning of *drA0334* in pET28a |
| 2 | pET28a-*dr2513* | CGGAATTCATGGCGTCCGCTTCTTCC (F)  CCGCTCGAGTCAGGGCACTTGCCTTTT (R) | Cloning of *dr2513* in pET28a |
| 3 | pET28a-*dr1243* | GGAATTCGTGCTGACCCTGCCTGCG (F)  CCGCTCGAGTCAGGCGTTCTCCCCCGC (R) | Cloning of *dr1243* in pET28a |
| 4 | pET28a-*drA0332* | CGGGATCCGTGACGGCCCCCCAGACC (F)  CCCAAGCTTCTAGATGCGCCGAATCAG (R) | Cloning of *drA0332* in pET28a |
| 7 | pET28a-*PP2C* | GGAATTCCGGGTGGCCGCCCGCACC(F)  CCGCTCGAGTCACGCCGTTCTCTCGATCACCAG (R) | Cloning of PP2C domain of *drA0334* in pET28a |
| 8 | pET28a-*D394A* | GGAATTCCATATGGTGAACGAGAACCACCCGCG (F)  CCGCTCGAGTCACGCCGTTCTCTCGATCACCAG (R) CGGCCAACGAAGCCTCCTACGGCTAC (SDM-F)  GTAGCCGTAGGAGGCTTCGTTGGCCG (SDM-R) | Generation of D394A mutant of drA0334 |
| 9 | pET28a-*D419A* | GGAATTCCATATGGTGAACGAGAACCACCCGCG (F)  CCGCTCGAGTCACGCCGTTCTCTCGATCACCAG (R)  CCTGCGTCTGCGCCGGCATGGGCGG (SDM-F)  CCGCCCATGCCGGCGCAGACGCAGG (SDM-R) | Generation of D419A mutant of drA0334 |
| 10 | pET28a-*drA0333* | GGAATTCCATATGATGAGCATCACCTGCGAAGT (F)  CCGCTCGAGTCAGTCCTGGTGGAAGGTGAGCA(R) | Cloning of *drA0333* in pET28a |
| 11 | pET28a-*Chimera* | GGAATTCCTGATGGTCGGGCGCTTC (F)  AGCGGCGCGGCGTTGAACACGCCGTTGG (Linker-R)  AACGCCGCGCCGCTGCCCATCTAC (Linker-F)  CCGCTCGAGTCACGCCGTTCTCTCGATCACCAG (R) | Cloning of chimera (FHA domain of DRA0333 and PP2C domain of DRA0334) in pET28a |
| 12 | pRAD-*dr1243* | CTGGGCCC GTGCTGACCCTGCCTGCG (F)  GCTCTAGATCAGGCGTTCTCCCCCGC (R) | Cloning of *dr1243* in pET28a |
| 13 | *drA0334* pKNT25/pUT18 | GCT CTA GAG GTGAACGAGAACCACCCG (F)  GG GG TAC CGG CGCCGTTCTCTCGATCAC (R) | Cloning of *drA0334* in pKNT25/pUT18 |
| 14 | *dr2513* pKNT25/pUT18 | GCT CTA GAG ATGGCGTCCGCTTCTTCC (XbaI)  GG GG TAC CGG GGG CAC TTG CCT TTT CAG (KpnI) | Cloning of *dr2513* in pKNT25/pUT18 |
| 15 | *rqkA* pKNT25/pUT18 | GCT CTA GAG ATGCCGCTGACCCCTGGA (XbaI)  GG GG TAC CGG CCCTTCCTGCTCGCTGCG (KpnI) | Cloning of *dr2518* in pKNT25/pUT18 |
| 16 | *dr1243* pKNT25/pUT18 | GCT CTA GAG GTGCTGACCCTGCCTGCG (XbaI)  GG GG TAC CGG GGCGTTCTCCCCCGCCGT (KpnI) | Cloning of *dr1243* in pKNT25/pUT18 |
